# HSP90α and KLK6 Co-Regulate Stress-Induced Prostate Cancer Cell Motility

**DOI:** 10.64898/2025.12.19.695259

**Authors:** Katelyn L. O’Neill, Johnny W. Zigmond, Raymond Bergan

## Abstract

**Background:** Prostate cancer (PCa) metastasis is reliant on the activity of proteases, such as matrix metalloproteinase-2 (MMP-2). While increased extracellular heat shock protein 90α (eHSP90α) has been linked to increased MMP-2 activity, this has not been examined in the context of cellular stress.

**Methods:** We examined stress-induced eHSP90α in human prostate cell lines by immunoblot. Fluorometric gelatin dequenching and zymography assays measured MMP activity. Wound healing and Matrigel drop invasion assays were used to quantify cell motility. HSP90α knockout (KO) cells were established with CRISPR/Cas9. Proteases were profiled with molecular inhibitors and protein arrays and validated by siRNA knockdown, immunoblot, and motility assays.

**Results:** Stress increased eHSP90 in 4 of 4 human prostate cell lines examined. Surprisingly, it concurrently decreased MMP-2 activity. The functional relevance of this was demonstrated when conditioned media from stressed cells decreased the motility of non-stressed cells. Screening for protease inhibitors that would rescue stress-induced decreases in MMP-2 activity identified a single serine protease inhibitor: aprotinin. Yet, rescue with aprotinin was lost in HSP90α KO cells. A protease array identified stress-induced increases in kallikrein-related peptidase 6 (KLK6). Knockdown of KLK6 rescued stress-induced MMP-2 activity and cell motility.

**Conclusion:** We identify a novel stress-induced extracellular network that regulates MMP-2 activity and cell motility. We identified KLK6 as a stress-induced extracellular protease leading to decreased MMP-2 activity and cellular invasion, while eHSP90α is required for the rescue of MMP-2 activity once KLK6 is neutralized.

## 1. Introduction

Cancer cells are exposed to many different stressors in their environment, such as nutrient deprivation, oxidative stress, hypoxia, thermal stress, ER stress, DNA damage, and immune stress [1, 2]. Multiple mechanisms allowing cancer cells to adapt to and resist various stressors have been identified and studied over the past several decades. These defenses often drive aggressive behaviors encompassed in the hallmarks of cancer, including increased proliferation, resistance to growth suppressors and apoptosis, immune evasion, and increased cell motility [3–6].

One of the most studied stress response proteins in cancer is heat shock protein 90 (HSP90). While initially studied in response to thermal stress, HSP90 is upregulated in response to many different forms of cellular stress [7–10]. Cytoplasmic isoforms include HSP90β and HSP90α. While HSP90β is constitutively expressed, HSP90α expression is stress-induced in normal cells and both constitutively expressed at higher levels as well as inducible in cancer cells. [11–13]. Intracellularly, HSP90β and HSP90α function as chaperones, employing different co-chaperones to assist in proper protein folding and re-folding as a means of resisting stress, protein damage, and apoptosis [7, 8, 14–16]. HSP90α has more recently been identified and studied as an extracellular protein (eHSP90α) that promotes migration and invasion in multiple types of cancer [17, 18]. It is proposed to do so by co-chaperone mediated activation of matrix metalloproteases (MMP), specifically MMP-2 and MMP-9 [19–24].

MMPs are extracellular proteins that break down extracellular matrix (ECM) components, enabling cells to be motile. This is especially important for cancer cells when invading out of the primary tumor site. MMPs have been extensively studied, and their regulation and function are complex, especially in cancer biology [25–27]. While multiple reports support that increases in eHSP90α increase MMP-2 expression and/or activity, none of these systems examine eHSP90α and MMP-2 in the context of cell stress [19–24]. This is relevant because cell stress is recognized as the physiological inducer of HSP90α, and prior reports either deployed engineered perturbations or were associative.

The first report of eHSP90α playing an extracellular role in migration and invasion through modulation of MMP-2 was over two decades ago [19]. In that study, investigators probed for cell surface proteins involved in invasion and identified cell surface and secreted HSP90α in fibrosarcoma and breast cancer (BrCa) cells. They looked at MMP-2 to explain effects on invasion and found that HSP90α interacted with and activated MMP-2 by utilizing immunoprecipitation assays and geldanamycin inhibition of HSP90α. Several studies have looked at the interaction of eHSP90α and MMP-2 in cancer cells by studying higher endogenous levels in aggressive cancer cell lines, exogenous gene expression, or adding recombinant protein to culture medium [21–24]. Many studies also employ eHSP90α inhibitors, such as antibodies, beads coated with HSP90 inhibitor geldanamycin or impermeable forms of geldanamycin to show its role in cell motility [19, 21, 22, 28, 29]. Importantly, none of these studies look at stress-induced eHSP90α associated with MMP-2 activity.

In the current study, we examine the impact of cell stress on eHSP90α, MMPs and cell motility in human prostate cancer (PCa). Cell motility is a fundamental cellular process whose dysregulation in cancer cells leads to initiation of metastatic behavior and invasion out of the primary tumor site [30]. Metastasis is the lethal manifestation of the vast majority of cancers. This is true in PCa, as the prognosis and survival for those with metastatic disease are extremely poor [31, 32]. Cell motility remains a difficult process to selectively target therapeutically. An improved understanding of how cell stress, a fundamental response pathway, regulates cell motility, a fundamental process central to cancer, will provide essential tools to inform optimization of patient risk stratification, tailored care and development of precision therapeutics.

This study sought to understand the impact of cell stress on MMP-2 activity and cell motility in human PCa, and the role of eHSP90α in regulating those processes. Here we identify a novel stress-induced extracellular network that regulates MMP-2 activity and cell motility. We demonstrate that cell stress, as induced by heat shock, increases eHSP90α but decreases MMP-2 activity in 4 of 4 human prostate cell lines and cell motility in 2 of 2 PCa cell lines examined. This finding is in contrast to reports in which artificially introduced eHSP90α increases MMP-2 and motility [22–24], which demonstrates the importance of examining stress-response pathways in the context of cell stress. We go on to show that stress-induced decreases in MMP-2 are rescued by adding the serine protease inhibitor, aprotinin. Yet, this rescue was lost in HSP90α KO cells. We then demonstrate that the serine protease, kallikrein-related peptidase 6 (KLK6), increased with cell stress and that its knockdown rescued stress-induced decreases in MMP-2 and cell motility. Together, these findings identify a stress-induced extracellular protease, KLK6, leading to decreased MMP-2 activation and cell motility, while demonstrating the requirement of eHSP90α for the rescue of MMP-2 activity once KLK6 is inhibited.

## 2. Materials and Methods

### 2.1. Cell culture

PC3 and LNCaP cells were obtained from the American Type Culture Collection (ATCC, Manassas, VA). PC3 and LNCaP cells were cultured in RPMI 1640 media (Gibco, no 18835030) supplemented with 10% FBS (Gibco, no. 16140-089 and 1% Antibiotic-Antimycotic (Gibco, no. 15240062). Normal and cancer cells from the same patient were micro-dissected and transformed yielding 1532NPTX (normal) and 1532CPTX (cancer), HPV-transformed primary cell lines were a generous gift from S. Topalian (National Cancer Institute, Bethesda, MD). 1532NPTX and 1532CPTX were cultured with keratinocyte-serum free medium, supplemented with recombinant epidermal growth factor (rEGF) and bovine pituitary extract (BPE) (Gibco) as well as 3% FBS and 1% Antibiotic-Antimycotic (Gibco, no. 15240062). All cells were maintained at 37°C in a humidified atmosphere of 5% carbon dioxide under sub-confluent exponential growth conditions and passaged ≥ 2 times weekly. All cell lines were drawn from stored stock cells, replenished on a standardized periodic basis, and were routinely monitored for Mycoplasma (Mycoplasma Detection Kit, Southern Biotech no.13100-01

All cell lines were authenticated and acquired through ATCC or the primary investigator, as above. Once acquired cells were cultured and expanded under quarantine conditions (i.e. a separate, dedicated incubator). They were stored as primary stocks and only used once confirmed to be negative for mycoplasma. Microscopy examination of morphology was conducted and required to match previously published phenotypes. All cell lines were handled one at a time and complete sterilization of working services was performed in between.

### 2.2. Antibodies and Reagents

Antibodies used: anti-GAPDH (Cell Signaling Technologies, no. 2118), anti-HSP90α (Cell Signaling Technologies, no. 8165), anti-HSP90β (Cell Signaling Technologies, no. 5087, Enzo Life Sciences, ADI-SPA-846-F), anti-KLK6 (R&D systems, no. AF2008) anti-MME (R&D systems, no. AF1182) anti-Rabbit IgG, HRP-conjugated (Cell Signaling Technologies, no. 7074), anti-Goat IgG, HRP-conjugated (R&D Systems, no. HAF017).

Reagents used: Matrigel (Corning, no. 354230), Halt Protease/Phosphatase Inhibitor cocktail (Thermo Scientific no. 78442), E-64 (Millipore Sigma, no. E3132), Aprotinin (MP Biomedicals, no. 0219115801), Bestatin (MP Biomedicals, no. 0215284401), Leupeptin (VWR International, LLC, no. 97063-234), Sodium Fluoride (Sigma Aldrich, S6776), Sodium Orthovanadate (Thermo Scientific Chemicals, no. J60191.AD), Sodium Pyrophosphate (Sigma Aldrich, no. S6422), B-glycerolphosphate (Sigma Aldrich, no. G9422) Gelatin B (Millipore Sigma, no. G9391), Bbs1 (New England Biolabs, no. R3539), Blasticidin S hydrochloride (Thermo Scientific Chemicals, no. AAJ67216XF)

### 2.3. Cell Stress Treatment

Stress was induced in cells by thermal shock. Cells were cultured in complete media and incubated for indicated heat shock (HS) times at 43°C in a humidified atmosphere of 5% carbon dioxide; control cells were incubated at 37°C under otherwise identical conditions. After HS, cells were washed 2x with PBS and then placed in serum free media (SFM). Cell fraction (CF) and conditioned media (CM) were collected 24 hours after HS for PC3 and LNCaP and 8- or 16-hours post HS for 1532CPTX. Serum-free conditioned media were spun down (10min, 1400xg) to remove debris and particles before use. Cells were washed 2x with PBS before harvesting to remove cross-contaminates from CM.

### 2.4. Western blot

Western blots were performed as described by us [33]. Briefly, cell fractions were lysed in RIPA buffer (Thermo Scientific, no. 89900) supplemented with Halt Protease/Phosphatase inhibitor cocktail (Thermo Scientific no. 78442) for 30min on ice. Following centrifugation at 16,000xg for 10 min at 4°C, supernatant was collected for the cell fraction. The protein concentrations of the cell lysates were measured using A280 measurements by Nanodrop 2000. CM was collected from cells at indicated times post treatment. Serum-free CM was first spun down (10min, 1400xg) to remove cell particles and debris and then concentrated via centrifugal filters (>10k, 4-15ml, Sigma Millipore Amicon UFC801024, UFC901024) for 25min at 3500xg. Sample concentrations were equalized based on CF measurements. Approximately 150 µg of total protein from each cell fraction along with equalized amounts of CM was resolved by SDS-PAGE and transferred onto a nitrocellulose membrane before incubation with primary and secondary antibodies for Western blot. Proteins of interest were detected by chemiluminescence with SuperSignal West Femto maximum sensitivity substrate (Thermo Scientific, no. 4096).

### 2.5. Gelatin Zymography

Zymography assays for measurement of MMP activity were performed as described previously [34, 35] with modifications. Briefly, equal amounts of concentrated conditioned media from cultured cells were mixed with 4x sample buffer (250 mM Tris, pH 6.8, 8% SDS, 40% Glycerol, 0.01% Bromophenol Blue) and incubated at 37°C for 20mins. Samples were separated on an 8% SDS polyacrylamide gel containing 0.1% gelatin. Gels were washed four times with zymography renaturing buffer (G Biosciences, no. 786-482) for 30 minutes each wash, followed by a 30-minute wash with distilled H2O and then incubated for 24-48hrs at 37°C in zymography developing buffer (G Biosciences, no. 786-481). After developing gels were stained with 0.1% Coomassie Brilliant Blue solution containing 10% acetic acid and 40% methanol for 60-90 minutes. Gels were destained with 10% acetic acid and 40% methanol solution and imaged for absence of staining. Images were captured with iBright Imaging System (Invtirogen) and inverted to see clear bands as opaque.

### 2.6. Wound healing assay

Wound healing assays were performed as described by us with modifications [36]. Briefly, assays used Imagelock plates (Satorius, BA-04856) and Incucyte® S3 imaging and analysis (Sartorius). PC3 cells were seeded at 1.2x10^5^ per well and 1532CPTX cells were seeded at 1x10^5^ per well. The following day cell monolayers were scratched, rinsed with 1x PBS and then overlaid with 5mg/ml Matrigel (Corning, no. 354230) and culture media. Plates were then placed in Incucyte® S3 and images were captured every 4 hours over the next 24-48 hours. Images were analyzed by Incucyte® S3 software and relative wound density, which accounts for the background density of the wound at the initial time point and changes in the cell density outside the wound region as well as in the wound area, was calculated and graphed over time using Prism 10 (Graphpad Software Inc).

### 2.7. Matrigel drop invasion assay

Drop invasion assays were performed according to previous reports with modification [37]. Trypsinized cells were resuspended in 5 mg/ml Matrigel (Corning, no. 354230) at 5x10^6^ cells/ml. Cells in Matrigel were plated as 5μl drops on 24-well culture dishes, allowed to solidify at 37°C for 15min and then culture media was added. Pictures of drops were taken every 24 hours using 2x objective on EVOS M5000 imaging station (Invitrogen). The area of cells that invaded out of the Matrigel was quantitated with ImageJ. Invaded area = (Area of drop + invaded cells) – (area of day 0 drop).

### 2.8. Proliferation assay

Cells were seeded at 15,000/well for PC3 and 10,000/well for 1532CPTX in a 12-well culture dish. One day post plating the Incucyte S3 was used to quantify cells/well every 8 hours for the next 72-96 hours. Fold change in cell number overtime was used to compare separate cell lines or treatments and graphed with Prism 10 (Graphpad Software Inc).

### 2.9. Gelatin Dequenching Assay

Gelatin dequenching (DQ) assays were performed with the EnzChek® Gelatinase/Collagenase Assay Kit (Invitrogen, no. E12055). The kit was used with centrifugal filtered CM according to the manufacturer’s instructions. In brief, CM was combined with 1x reaction buffer and 100µg/ml Gelatin Fluorescein Conjugate and desired inhibitors in a black 96-well plate with a clear bottom (Corning, no. 3603). Negative controls with buffer -/+ each inhibitor and serum-free culture media -/+ inhibitors as well as positive controls with 1ng of rhMMP-2 (Abcam, no. ab81550) -/+ each inhibitor were run in tandem on each plate. Plates were read on a BioTek Synergy plate reader at 485/20 excitation and 528/20 emission with measurements every 2 hours over a 24-hour (hr) period. After subtraction of background, arbitrary fluorescence units for each sample were graphed and analyzed with Prism 10 (Graphpad Software Inc).

### 2.10. Protease and Phosphatase inhibition

Protease and phosphatase inhibitors were added to CM after centrifugal filtration. Halt Protease/Phosphatase inhibitor cocktail (Thermo Scientific no. 78442) was added 1:100 in CM. Protease inhibitors were added at the following final concentrations: 10 μM Bestatin, 1 μg/ml Aprotinin, 10 μM Leupeptin, and 15 μM E-64. Phosphatase inhibitors were added at the following final concentrations: 50 mM sodium fluoride, 1 mM sodium orthovanadate, 5 mM sodium pyrophosphate and 40 mM β-glycerophosphate. After addition of inhibitors samples were incubated at 37°C for 20min before running gelatin zymography. Samples for gelatin DQ assays were added directly to plates and then combined with Gelatin Fluorescein Conjugate before taking fluorometric measurements.

### 2.11. CRISPR constructs and knockout

Integrated DNA Technologies CRISPR design tool was used to design a pair of custom DNA oligos for the human *HSP90AA1* (GeneID: 3320) sgRNA targeting site (https://www.idtdna.com/site/order/designtool/index/CRISPR_PREDESIGN). Oligos were annealed and ligated into Bbs1-cut pSpCas9(BB)-2A-Blast (this plasmid was a gift from Ken-Ichi Takemaru; Addgene plasmid # 118055 ; http://n2t.net/addgene:118055). Oligo sequences were F-caccgACTTTTGTTCCACGACCCAT, R-aaacATGGGTCGTGGAACAAAAGTc. Successful cloning was verified with Sanger sequencing (Azenta, Genewiz) and hU6 Forward sequencing primer-GAGGGCCTATTTCCCATGATT. FugeneHD reagent (Avantor, no. MSPP-HD-1000) was used for transfection in PCa cells according to the manufacturer’s instructions. For CRISPR transfection 2.5 × 10^5^ cells were seeded in 6-well culture dish 24 hours prior to transfection. Four hundred nanograms of CRISPR construct was co-transfected with 100 ng of the corresponding pEGFP-N1 reporter plasmid. pcDNA3.1 was added to reach a total DNA concentration of 1.5 µg per transfection. The transfected cells were split 1:2 24hr later followed by 5 ug/ mL Blasticidin S hydrochloride (Thermo Scientific Chemicals, no. AAJ67216XF) selection for 72 hours. After selection cells were plated for single clone isolation in 96-well plates.

### 2.12. Genomic sequence analysis

Genomic DNA was isolated from PCa cells using DNeasy blood and tissue kit (Qiagen, no. 69504) and subjected to polymerase-chain reaction (PCR) to amplify genomic target regions of interest using Phusion polymerase (New England Biolabs, #E0553S). PCR was performed according to the manufacturer’s instructions and thermocycler settings were adjusted according to product size and optimal primer annealing temperatures. Purified PCR products were then sent for Sanger and/or Amplicon-EZ sequencing using primers from the original PCR to directly sequence (Azenta, Genewiz). Genomic amplicons were obtained using the following primers for *HSP90AA1* (Gene ID 3320) CRISPR target area:

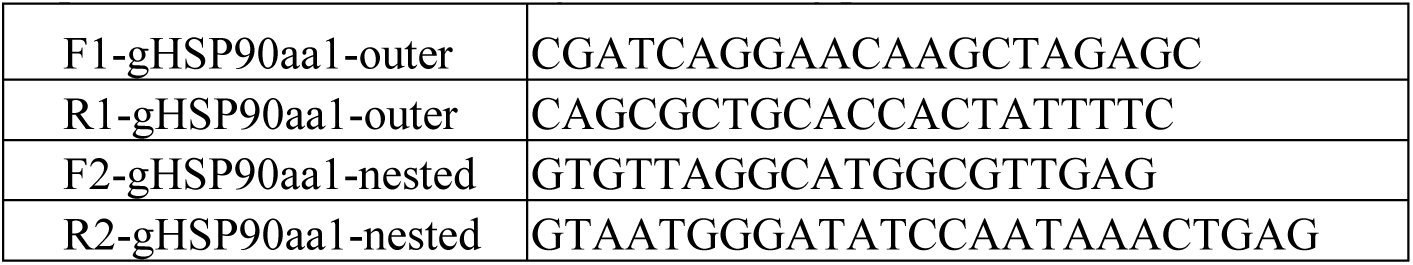

### 2.13. Migration assay

Cell migration assays were performed as described by us [38]. Briefly, cells were seeded in Boyden chamber 24-well plates (Corning, no 3422) at 50,000 cells/well. Chambers with cells contained media or conditioned media with 1% FBS and reservoirs below contained media or conditioned media with 10% FBS. Cells were allowed to migrate for 24 hours. Following this they were fixed and stained for DAPI (4′,6-diamidino-2-phenylindole, BD Biosciences, 564907) and migrated cells on the bottom of each chamber were quantified with three 10x images per well, taken by EVOS M5000 imaging station.

### 2.14. Protein Array

Protein arrays were performed with R&D systems Proteome Profiler^TM^ Array for human proteases (no. ARY025) according to manufacturer instructions. Membranes were blocked with array buffer for 1 hour. Centrifugal-filtered CM from untreated and stressed cells were incubated with biotinylated protease detection antibody cocktail for 1 hour and then added to prepared membranes. After overnight incubation at 4°C, membranes were washed with wash buffer and then incubated with Streptavidin-HRP for 30 minutes rocking at room temperature. Membranes were washed a second time and then prepared for development with provided Chemi Reagent Mix. Invitrogen iBright Imaging System (Invitrogen) was used to detect chemiluminescence. Pixel densities for each spot were measured with ImageJ software and normalized to reference spots and negative controls.

### 2.15. siRNA transfections

Knockdown using siRNA was performed as previously described by us [39] with modifications. Transfections were performed on 2.5×10^5^ cells in 6-well culture dishes with antibiotic-free RPMI medium supplemented with 10% FBS. DharmaFECT2 transfection reagent (Dharmacon, Inc., no. T-2002-01) was used for PC3 cells and DharmaFECT4 (T-2004-01) was used for 1532CPTX cells, per manufacturer’s instructions. On-TargetPlus Smart pools for siKLK-6, siMME, or siControl (Dharmacon, Inc., nos. L-005917-00-0005, L-005112-00-0005, D-001810-10-05) were suspended in 1× siRNA buffer (Dharmacon, Inc., no. B-002000-UB-100) and added to cells at a final concentration of 20 nM. On-TargetPlus Smart pools contain 4 different targeting siRNA for the same gene. Transfected cells were subjected to HS at 48hr after transfection and harvested 72hr after transfection for Western blot analysis or CM treatments for Matrigel drops or proliferation assays.

### 2.16. Statistical analysis

Data analysis was performed with GraphPad Prism Statistical Software version 10. Statistical comparisons between two groups were evaluated with the Welch’s Student’s t-test. One-way ANOVA was used to compare 3 or more samples in bar graphs. Two-way ANOVA (mixed model) was used to determine significance on XY graphs. P-values ≤0.05 are considered significant. The values were presented as the means ± SD in bar graphs and means ± SEM in XY graphs.

## 3. Results

### 3.1. Effects of cellular stress on HSP90 and MMP-2 in prostate cancer

Initially, we assessed the role of eHSP90α in stress-induced cell motility, as it has been linked to increased invasion in several types of cancers cells including breast, fibrosarcoma, melanoma and PCa [19, 22, 24, 29, 40]. Previous reports examined models in which HSP90α was artificially introduced, inclusive of overexpression by lentivirus or adding recombinant HSP90α to culture media [22–24, 40]. We reasoned that inducing expression by applying cellular stress would be more physiologically relevant. Multiple PCa cell lines were examined: PC3 is metastatic hormone refractory androgen receptor (AR) negative [41], LNCaP is metastatic AR positive [42], while 1532NPTX and 1523CPTX are paired primary normal and cancer cells, respectively, isolated from the same patient [43]. Stress was induced by heat shock, after which cell fraction and conditioned media were collected separately to look at intracellular versus extracellular protein changes. All four cell lines examined displayed an increase of both extracellular HSP90α (eHSP90α) and eHSP90β when subjected to stress (Figure 1A-C). PC3, LNCaP and 1532NPTX cells exhibited an increase of intracellular HSP90α (iHSP90α) with stress, consistent with iHSP90α known to be inducible, while iHSP90β levels remained relatively constant for PC3, 1532NPTX and 1523CPTX, but decreased in LNCaP cells, consistent with iHSP90β known to be constitutively expressed. Together, these findings demonstrate that eHSP90α and eHSP90β increase with cell stress and this is observed in four of four prostate cell lines examined.

**Figure 1.**
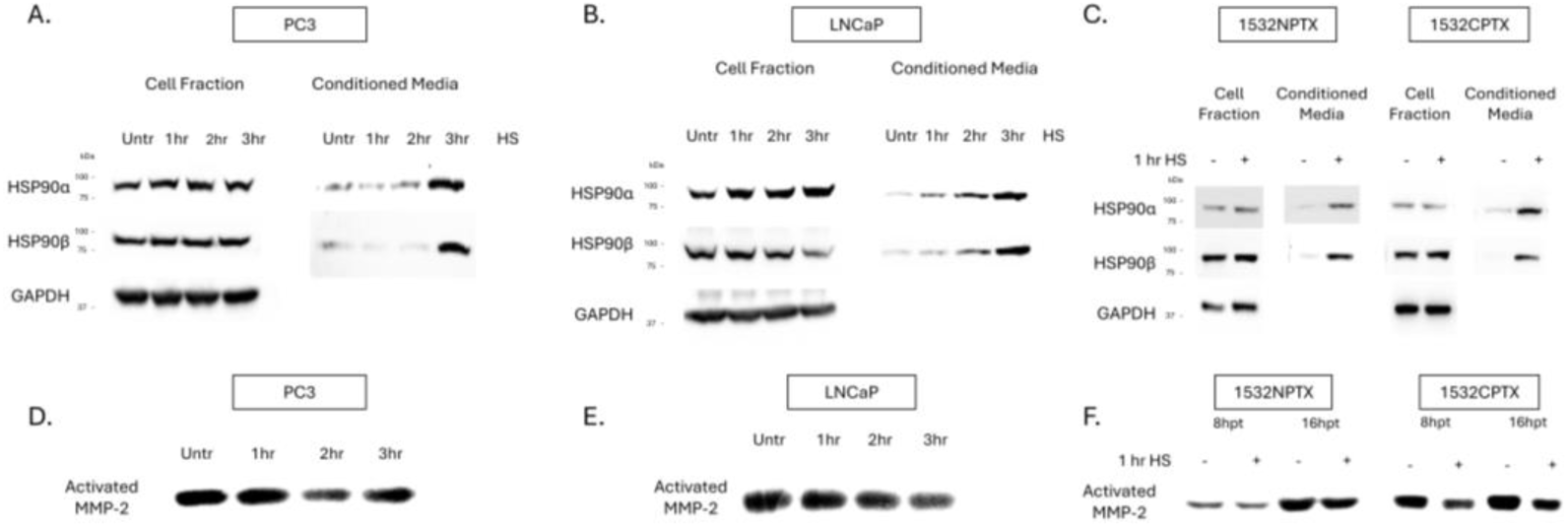
Stress-induced protein changes in PCa cells lead to increased eHSP90 and decreased MMP-2 activity. (**A-C**) Stress-induced heat shock protein 90 (HSP90) expression and secretion in established metastatic prostate cancer (PCa) (PC3, LNCaP) and paired primary (1532NPTX and 1532CPTX) cells. Cells were heat shocked (HS, 43°C) for indicated times and then place in serum-free media (SFM). Cell fractions and conditioned media were collected and analyzed 24-hours (hr) post heat shock in PC3 and LNCaP and 8hr post heat shock for primary cells (n= 2-3). (**D-F**) Matrix metalloproteinase-2 (MMP-2) activity in conditioned media (CM) from PC3, LNCaP, and 1532NPTX and 1532CTPX cells as indicated by gelatin zymography n=3. CM as collected and analyzed 24hr post stress in PC3 and LNCaP and 8hr and 16hr post stress for primary cells (n= 2-3). Images are inverted. Bands are indicative of gelatin degradation due to MMP-2 activity.

While prior studies report that increases in eHSP90α increase MMP-2 activation, they do so either through artificial alterations in eHSP90α, or through comparisons between high versus low eHSP90α cell lines [22–24]. Importantly, none of these reports examine the impact of stress-induced eHSP90α. In apparent contrast to those reports, it can be seen in Figure 1D-F that in the context of stress, increased eHSP90α is associated with decreased MMP-2 activation. Of importance, this was seen in four of four different human prostate cell lines examined. MMP activity depicted in Figure 1 was measured by gelatin zymography, which detects MMP-2 and MMP-9 activity [44]. Of importance, MMP-9 activity is much lower than MMP-2 (Figure S1). Further, MMP-9 was minimally impacted by stress in PC3 and 1532NPTX cells, was decreased in LNCaP cells and exhibited a biphasic increase then decrease with length of time post stress in 1532CPTX cells. Together, these findings demonstrate that under conditions of cell stress, increased eHSP90α is associated with decreased MMP-2, this effect is observed across multiple cell lines, MMP-9 activity is much lower than MMP-2, and it exhibits inconsistent effects across cell lines and over time.

### 3.2. Effects of extracellular stress response proteins on PCa cell invasion

We next sought to examine the mechanisms of these effects. This is important for several reasons. Our findings reveal different results from prior reports. Other reports have either not examined underlying mechanisms or have done so only under conditions of engineered expression and/or lack of cell stress with model systems [19–24]. Also, we focused our studies on MMP-2. This is because MMP-2 is recognized as a major MMP whose expression and activity has been closely tied to cell invasion and the metastatic phenotype across a wide range of cancer types, inclusive of PCa [34, 35, 45–54]. In addition to the proven biological importance of MMP-2, its uniform response to stress across multiple cell lines in the current study, its activity being much higher than MMP-9, and MMP-9 exhibiting inconsistent responses, further support focusing on MMP-2.

We first assessed if the observed decrease in MMP-2 activity had an impact on cell motility. Further studies focused on PC3 and 1532CPTX cells, providing representation of a metastatic variant and primary PCa cells, respectively. CM from control (non-stressed) cells and those subjected to stress were collected and then applied to naive cells in the context of wound healing or Matrigel drop invasion assays. As can be seen in Figure 2A-D and Figure S2A-D, CM from stressed cells significantly decreased invasion compared to CM from control cells and did so in both PC3 and 1532CPTX cells. For wound healing assays, significant effects were evident 8 and 12 hr and persisted until 24 and 36 hr experimental termination points for PC3 and 1532CPTX cells, respectively (Figure 2A, B, Figure S2A, B). For Matrigel drop invasion assays, significant effects were evident at 3 days for both cell types and persisted until 5 and 4 day termination points for PC3 and 1532CPTX cells, respectively (Figure 2C, D, Figure S2C, D). To ensure that invasion assays were not influenced by different cell growth rates, cell growth assays were conducted and demonstrated no significant difference between control and stress CM treatments (Figure 2E, F).

**Figure 2.**
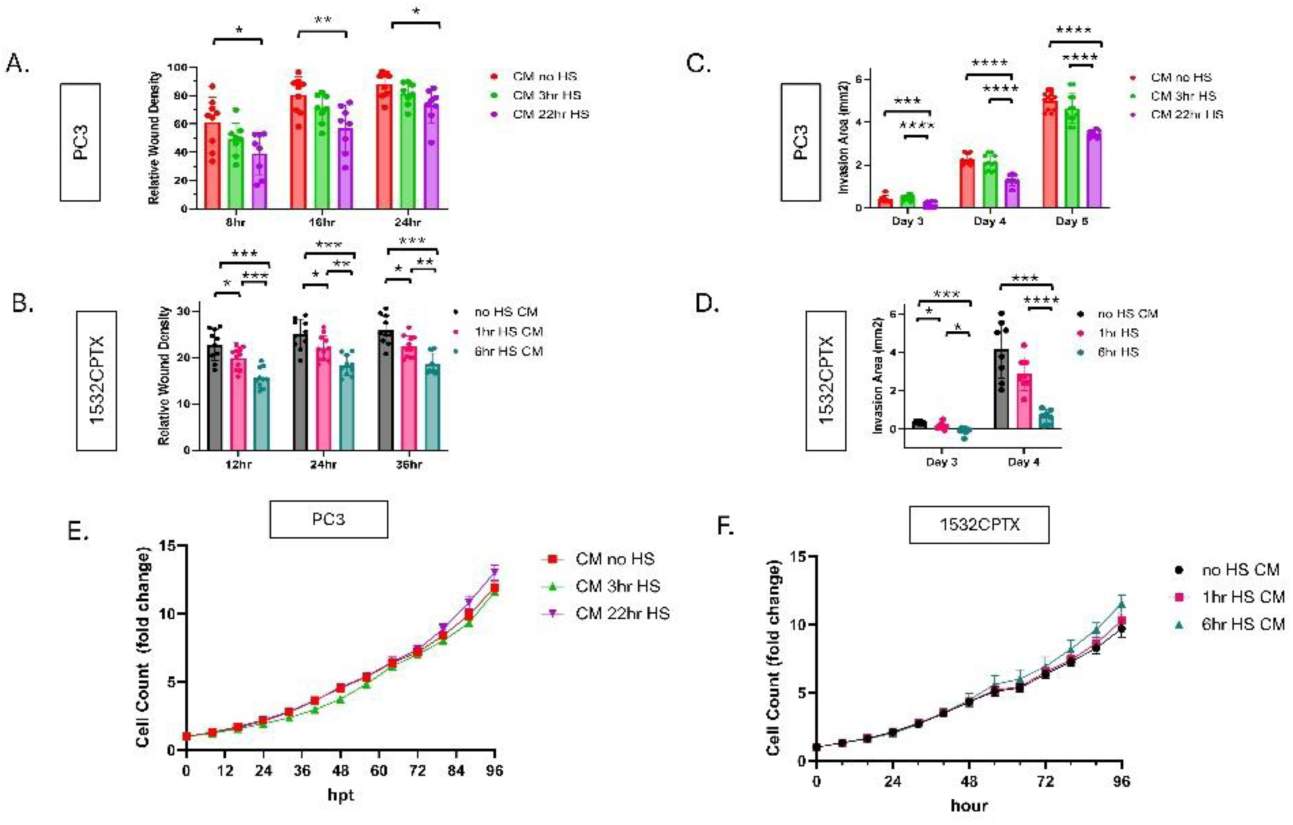
Stress-induced extracellular proteins lead to decreased cellular invasion. (**A, B**) Wound healing invasion assay for untreated PC3 and 1532CPTX cells incubated with CM from untreated and stressed cells). Cell monolayers were scratched and then overlaid with Matrigel and treated with CM from cells -/+ stress. PC3 CM was harvested 24 hours post stress and 1532CPTX CM was harvested 16hr post stress. Relative wound density was measured by Incucyte S3 live cell analysis over 24-36hrs. (**C, D**) Matrigel drop invasion assay with PC3 and 1532CPTX cells treated with CM from untreated and stressed cells. (**E, F**) Proliferation assays with PC3 and 1532CPTX treated with CM from -/+ stress samples. Two to four individual biological replicates were performed for the above experiments, and the results were combined. Column graph error bars indicate SD, XY graph error bars are SEM, * p< 0.05, ** p< 0.01, *** p<0.001, **** p<0.0001. Welch’s t-test was used to determine statistical significance.

These findings demonstrate that CM from stressed cells decreases cell motility. Their concordant findings with decreased MMP-2 in stressed cells supports the physiological relevance of MMP-2 in the current system. Further, findings align with the established role and importance of MMP-2 in regulating motility. Together, these results indicate that cellular stress results in a novel extracellular regulatory mechanism that decreases MMP-2 activity, resulting in decreased cellular invasion in the context of higher levels of eHSP90α.

### 3.3. Influence of other stress-induced factors on the regulation of MMP-2 activity

We hypothesized that a stress-induced protease or phosphatase in CM was leading to the decrease in observed MMP-2 activity. Proteases and phosphatases are known regulators of proteins and they both have been reported to impact MMP activity [55–62]. To begin examining this concept, CM was collected from untreated and stressed cells, a combination of protease/phosphatase inhibitors (PPIs) was added, and MMP-2 activity was measured in PC3 and 1532CPTX samples. As shown in Figure 3A and B, PPI rescued MMP-2 activity in both cell lines examined. PPI did not exhibit a rescue effect on MMP-9 in either PC3 or 1532CPTX cells (Figure S3).

**Figure 3.**
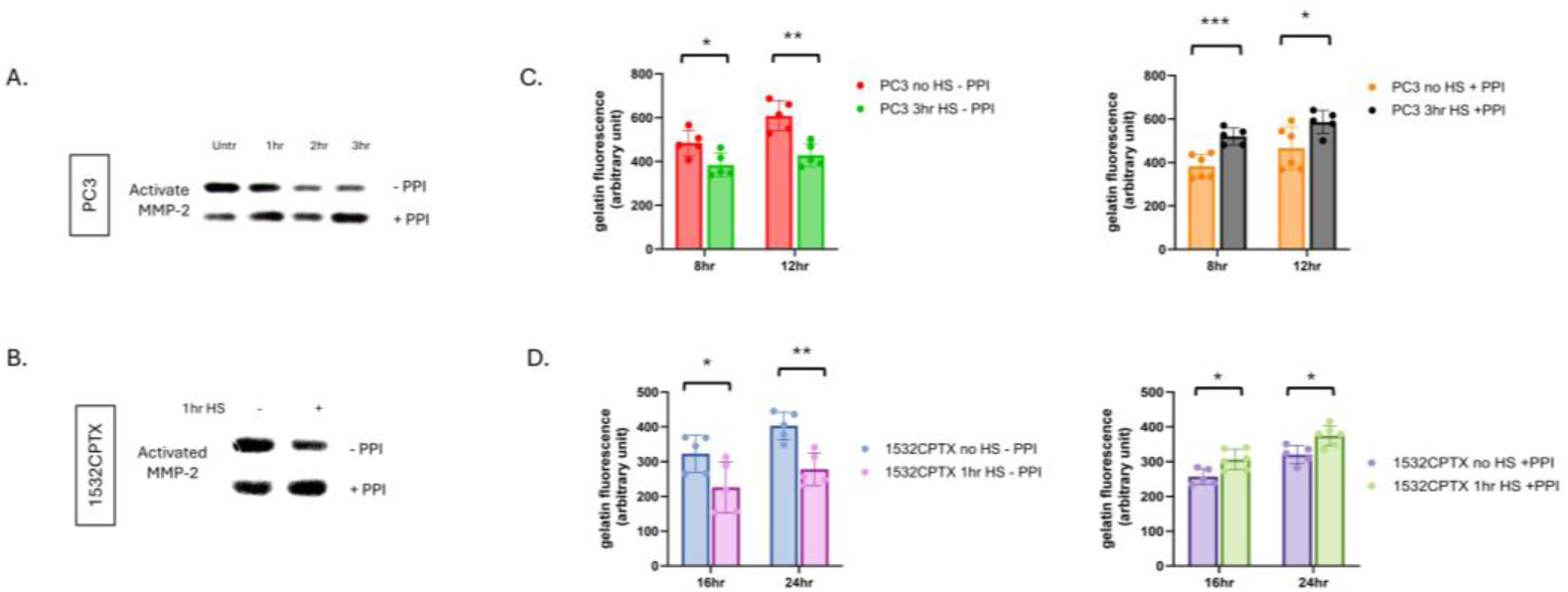
Protease and phosphatase inhibition results in increased stress-induced MMP-2 activity. (**A**) MMP-2 activity in CM from PC3 cells 24hr post stress and (**B**) 1532CPTX cells 8hr post stress -/+ protease/phosphatase inhibitors (PPI) as indicated by gelatin zymography n=2-3. (**C**) MMP-2 activity in CM from PC3 24hr post stress -/+ PPI and (**D**) 1532CPTX CM 16hr post stress -/+PPI as measured by gelatin dequenching (DQ) assays. Graphs represent combined results from at least 2 biologically independent experiments. Error bars indicate SD, * p< 0.05, ** p< 0.01, *** p<0.001. Welch’s student t-test was used to determine statistical significance.

Gelatin dequenching (DQ) assays offer an orthogonal measure of gelatinase activity. Compared to gelatin zymography, they represent a newer and more robust methodology, provide a quantitative output of activity and readily support multiple analyses. Like zymography, gelatin DQ assays measure the activity of MMP-2 and MMP-9, the predominant gelatinases in epithelial cells [63, 64]. In the context of the current models, MMP-9 activity is minimal and thus gelatin DQ assays provide a measure of MMP-2 activity. Gelatin DQ assays showed that CM from stressed cells exhibited significantly decreased gelatinase activity compared to control cells, and this was seen in both PC3 and1532CPTX cells (Figure 3C, D). Further, PPI rescued gelatinase activity in both cells examined. Findings in gelatin DQ assays emulate those in zymography. Together, these findings demonstrate that stress-induced decreases in MMP-2 activity can be rescued and that a protease and/or phosphatase is responsible, directly supporting our hypothesis.

### 3.4 Effects of eHSP90α on MMP-2 activity

To examine the role of eHSP90α in stress-induced changes in MMP-2 and cell motility, we established the first reported PCa HSP90α knockout (KO) cell line utilizing CRISPR/Cas-9 targeted gene editing [65]. For this study, a CRISPR-expression vector targeting *HSP90AA1* was transfected in PC3 cells. Blasticidin-selected pools were analyzed for loss of expression via Western blot (Figure S4A) and plated for isolation of single clones. Individual clones with successful KO were verified with immunoblot (Figure 4A) and genomic sequencing (Figure S4B). Empty vector (EV) control clones were selected and isolated in parallel. Loss of HSP90α did not influence proliferation rates, with EV and KO clones having similar doubling times as parental PC3 cells (Figure 4B). Of importance, CM from HSP90α KO cells exhibited a strong increase in MMP-2 activity for both gelatin zymography and gelatin DQ assays, compared to CM from EV clones (Figure 4C, D). Given the decrease in MMP-2 observed after stress-induced increases in eHSP90α, this increase in MMP-2 with HSP90α KO provides complementary orthogonal findings. These findings demonstrate that it is possible to successfully knockout HSP90α in PCa cells and eHSP90α is playing a role in regulating MMP-2 activity.

**Figure 4.**
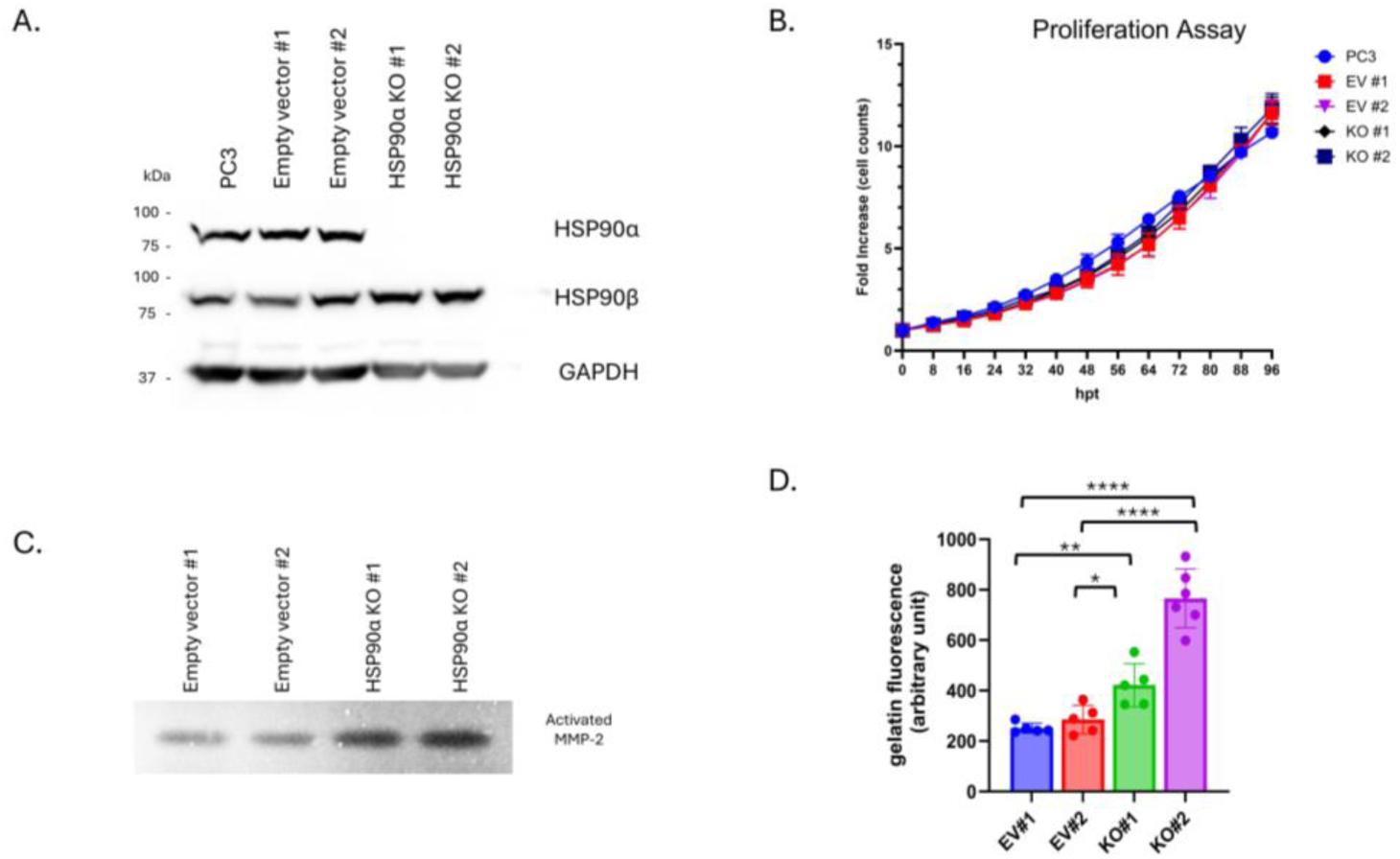
HSP90α KO cells exhibit increased MMP-2 activity. (**A**) Western of PC3 parental cell line, empty vector (EV) control single clones #1 and #2 and HSP90α KO single clones #1 and #2. (**B**) Proliferation assay of PC3, empty vector and HSP90α KO clones. (**C, D**) Zymography and gelatin DQ assay showing MMP activity in CM from HSP90α KO clones and empty vector controls. Graphs represent combined results from at least 2 biologically independent experiments. Bar graph error bars indicate SD, XY graph error bars are SEM. Statistical significance was determined by Welch’s student t-test, * p<0.05, **<0.01, ***<0.001, ****<0.0001.

Ironically, while KO cells display increased extracellular MMP-2 activity, the cells themselves have decreased motility compared to EV controls when assayed for both migration and invasion (Figure S5 A-C). Collectively, these experiments demonstrate that multiple PC3 HSP90α KO clones exhibit a similar phenotype characterized by increased extracellular MMP activity and decreased migration and invasion, but with unaltered cell growth.

Additionally, 1532CPTX HSP90α KO clones were established alongside 1532CPTX EV controls. Knockout was verified via immunoblot and genomic sequence analysis (Figure S6A, B). Increased MMP-2 activity was observed in 1532CPTX HSP90α KO CM compared to EV controls via gelatin DQ assays (Figure S6E). However, decreased cell motility was also observed in 1532CPTX KO clones compared to EV clones as was decreased proliferation (Figure S6 C, D). As decreased cell growth would confound further assays and analyses, 1532CPTX KOs were not used in subsequent studies.

Next, we examined the impact of cellular stress on HSP90β induction and MMP-2 activity in HSP90α KO cells. Cellular stress in KO cells led to increases in cytosolic and extracellular HSP90β, presumably representing a compensatory mechanism in the face of loss of HSP90α (Figure 5A, Figure S7A). In control EV cells, there was no change in cytosolic HSP90β with stress, but there was induction of both cytosolic and extracellular HSP90α, as seen in parental cells (Figure 5A).

**Figure 5.**
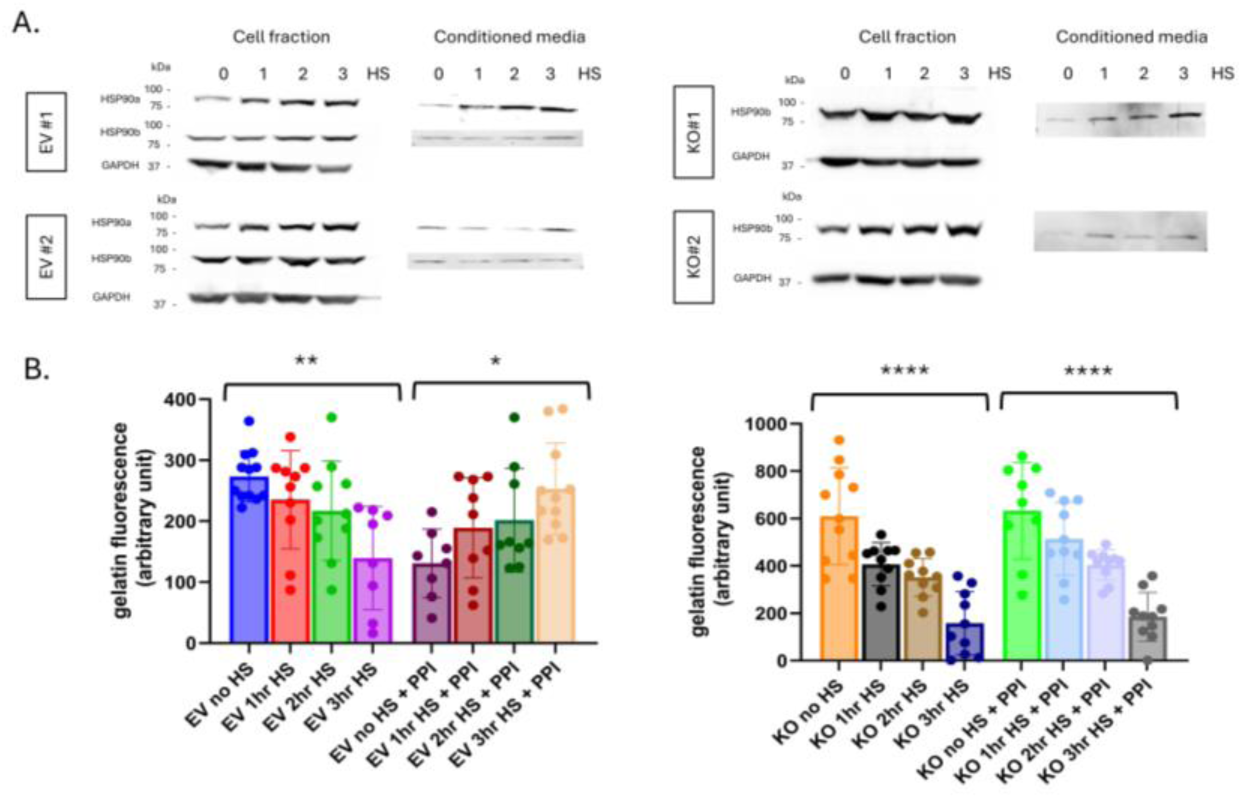
Effects of cellular stress in HSP90α KO cells. (**A**) Stress-induced HSP90α and/or HSP90β expression in cell fractions and conditioned media from empty vector controls and HSP90α KO PC3 cells. CM was harvested 24hr post stress. (**B**) MMP-2 activity in CM from untreated and stress treated samples as measured by gelatin DQ assay -/+ PPI. Three individual biological replicates were performed for the above experiments, and the results were combined. Statistical significance was determined by one-way ANOVA, error bars indicate SD, * p<0.05, **<0.01, ***<0.001, ****<0.0001.

Subsequently, MMP-2 activity was examined in KO and EV cells under cell stress, and by gelatin DQ to support comparison and quantification (Figure 5B). MMP-2 activity is almost two times higher in untreated KO cells compared to untreated EV cells, again demonstrating that HSP90α plays a role in regulating MMP-2 under non-stress conditions. Despite the higher MMP-2 activity in untreated KO cells, after stress MMP-2 activity declined to nearly identical levels in KO and EV cells. This demonstrates that HSP90α is not necessary for stress-induced decrease of extracellular MMP-2 activity. Further, and of high interest, while PPI rescued stress-induced decrease of MMP-2 activity in EV cells, it failed to do so in KO cells (Figure 5B). This demonstrates that HSP90α is necessary for PPI-mediated rescue of MMP-2 activity. That is, HSP90α is necessary for the provision of factors that are the targets of PPI in this system. These findings demonstrate that HSP90α plays a role in the regulation of MMP-2 activation. It lowers MMP-2 activity nder non-stress conditions. It is not, however, required for the decreased MMP-2 activity induced by cellular stress. Conversely, HSP90α is required for PPI-mediated rescue of stress-induced increase in MMP-2.

### 3.5. The Role of Proteases and Phosphatases in regulating MMP-2 activity

Having demonstrated that PPI affects the regulation of MMP-2, investigations next sought to examine this further with the goal of identifying which factors contribute to MMP-2 regulation. The PPI mixture contained four protease inhibitors: Bestatin, Aprotinin, Leupeptin, and E-64 and four phosphatase inhibitors: sodium fluoride, sodium orthovanadate, sodium pyrophosphate and β-glycerophosphate. To determine whether a protease or phosphatase was responsible for stress-induced decrease in MMP-2 activity, we pooled the protease inhibitors together and the phosphatase inhibitors together, then added them to CM from PC3 or 1532CPTX cells, comparing cells subjected to stress to controls (Figure 6A-D). Rescue was evident in both PC3 and 1532CPTX cells with addition of protease inhibitors. However, with phosphatase inhibitors rescue was not present in 1532CPTX cells nor PC3 cells at the 3hr time point but was preceded by a biphasic response at earlier time points. These findings provided strong support that a stress-induced protease was responsible for the decreased MMP-2 activity.

**Figure 6.**
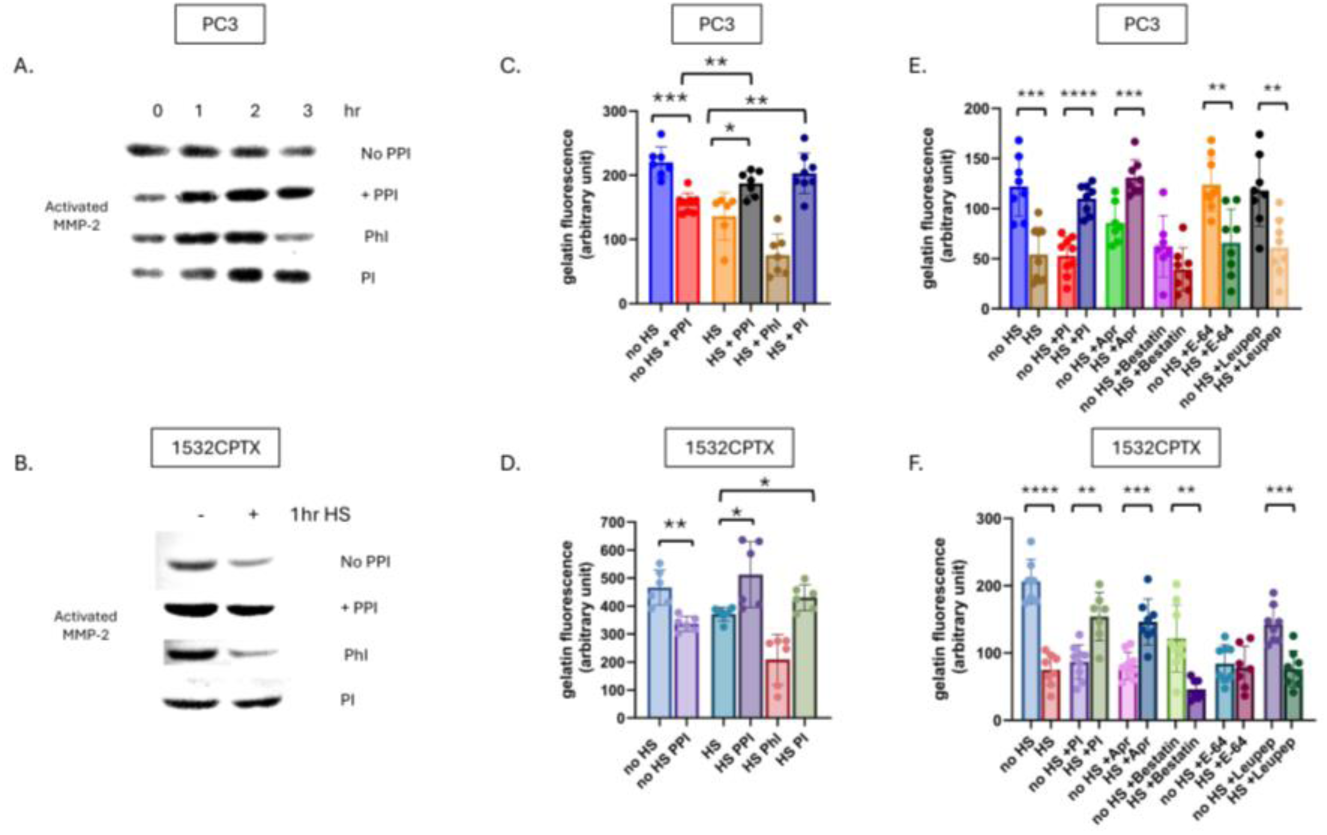
Stress-induced protease activity leads to decreased MMP-2 activity. (**A**) MMP-2 activity in PC3 CM 24hr post stress with PPI, phosphatase inhibitors (PhI: sodium fluoride, sodium orthovanadate, sodium pyrophosphate and β-glycerophosphate) or protease inhibitors (PI: Bestatin, Aprotinin, Leupeptin, and E-64) as indicated by gelatin zymography, n=2. (**B**) Gelatin zymography with 1532CPTX CM 8hr post treatment from untreated and stressed cell with PPI, PhI and PI n=2. (**C**) MMP activity as measured by gelatin DQ assay in PC3 CM 24hr post stress and (**D**) 1532CPTX CM 16hr post stress with PPI, phosphatase inhibitors or protease inhibitors added. (**E**) Gelatin DQ assay with individual protease inhibitors: Bestatin, Aprotinin, Leupeptin, and E-64 added to PC3 HS CM 24hr post stress; and F) 1532CPTX CM 16hr post stress. Graphs represent combined results from 3 biologically independent experiments. Statistical significance was determined by Welch’s student t-test, * p<0.05, **<0.01, ***<0.001, ****<0.0001.

Two independent and complementary approaches were pursued to determine which protease(s) were involved in decreased MMP activity. In one approach, individual protease inhibitors were added to CM from untreated and stressed PC3 and 1532CPTX cells, and resultant MMP-2 activity was measured (Figure 6E and F). Across both PC3 and 1532CPTX cells the only protease inhibitor that rescued MMP-2 activity was the serine protease inhibitor, aprotinin. The second approach screened for proteases in CM whose expression increased after cell stress. We hypothesized that the responsible protease would be increased upon cellular stress and subsequently lead to decreased MMP-2 activity. The Proteome Profiler™ Human Protease Array (R&D systems) supports detection and semi-quantification of 35 different proteases (Figure 7A and B). Using it to compare control and stressed CM from PC3 and 1532CPTX cells, two proteases were found to be significantly increased with stress in both cell lines: Kallikrein-related peptidase 6 (KLK6) and Membrane metallo-endopeptidase (MME/CD10). Increased levels of both proteases in CM after cell stress were verified via Western blot (Figure 7C and D). KLK6 is strongly inhibited by aprotinin [66, 67]. Prior reports indicate that MME is not significantly inhibited by protease inhibitors examined in the current study [68–74].

**Figure 7.**
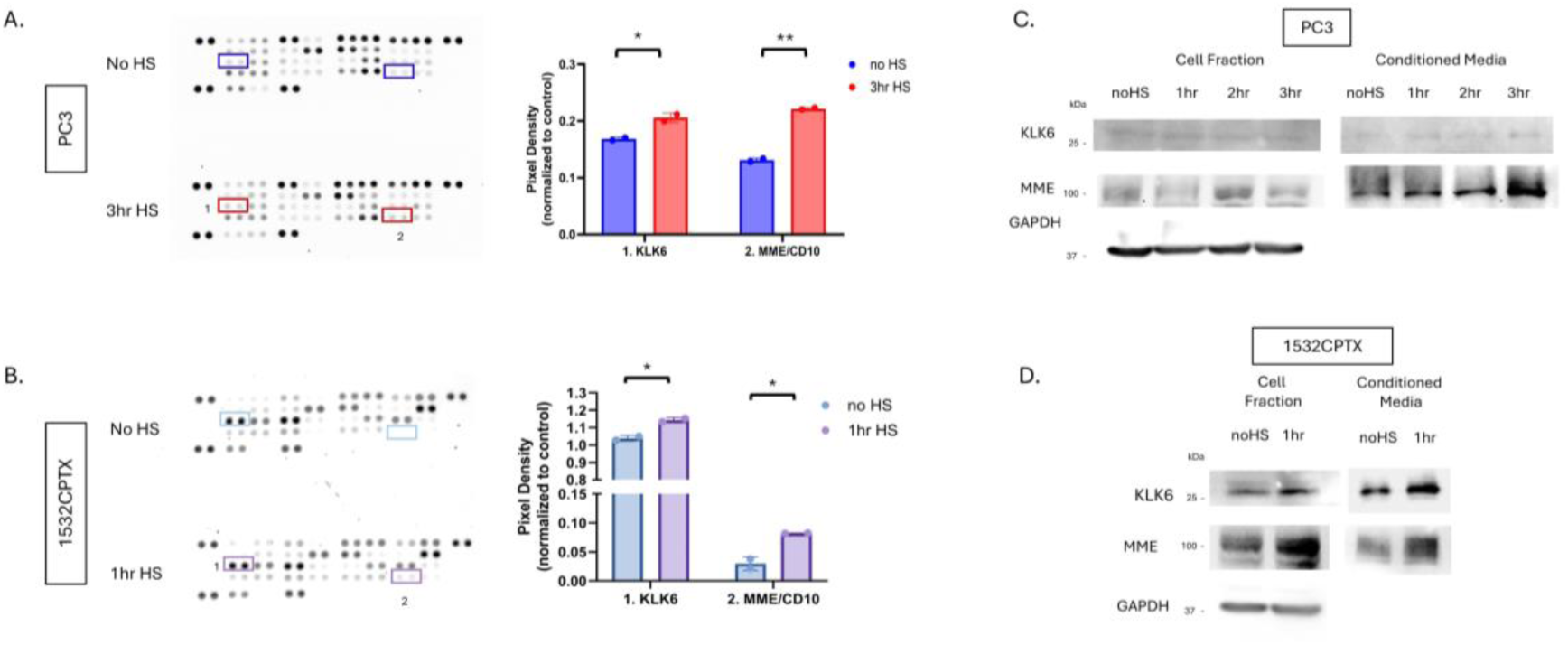
MME and KLK6 proteases are increased in conditioned media in a stress-dependent manner. Protease profiler array membranes (R&D systems) with 35 protease capture antibodies printed in duplicate were probed with CM from (**A**) PC3 24hr post stress and (**B**) 1532CPTX cells 16hr post stress. Pixel density after immunoblot exposure was used to identify common proteases between the two cell lines that were significantly increased (red/purple boxes): Kallikrein-related peptidase 6 (KLK6) and Membrane metallo-endopeptidase (MME/CD10). (**C, D**) Immunoblot assays were used to detect KLK6 and MME -/+ stress in PC3 and 1532CPTX cell fraction and CM n=3. Statistical significance was determined by unpaired student t-test, * p<0.05, **<0.01.

Together, these findings identify protease activity as mediating stress-induced decreases in MMP-2 activity. They further indicate that the function of a serine protease is responsible. An analysis of changes in protein expression identified KLK6 and MME as candidate mediators of this function. Considering protease inhibitors used previously would have little to no effect on MME, KLK6 emerged as the priority candidate.

### 3.6. Proteases involved in regulation of MMP activity and subsequent cellular invasion

To determine the role of MME and KLK6 in PCa cell motility, we performed siRNA knockdown. We hypothesized that knockdown of either KLK6 or MME would mimic the effect of PPI in CM. MME or KLK6 specific siRNAs were transfected into PC3 and 1532CPTX cells, non-targeting negative control siRNAs were transfected in parallel. Cells were subjected to stress 48hrs post transfection and the cell fractions and CM were harvested 24hr later for PC3 cells and 16hr later for 1532CPTX. As can be seen, KLK6 and MME were readily detected by Western blot for CM, and their successful knockdown was evident for PC3 and 1532CPTX cells, both in untreated and stressed cells (Figure 8A, Figure S8A). Of note, PC3 also maintained stress-induced increases of MME and KLK6 in control samples (Figure 8A). 1532CPTX cells did not exhibit increases in MME or KLK6 when subjected to stress post siRNA transfection (Figure S8A). This may be attributed to stress induced by transfection alone. Due to loss of this increase, we focused on PC3 cells for further functional assays.

**Figure 8.**
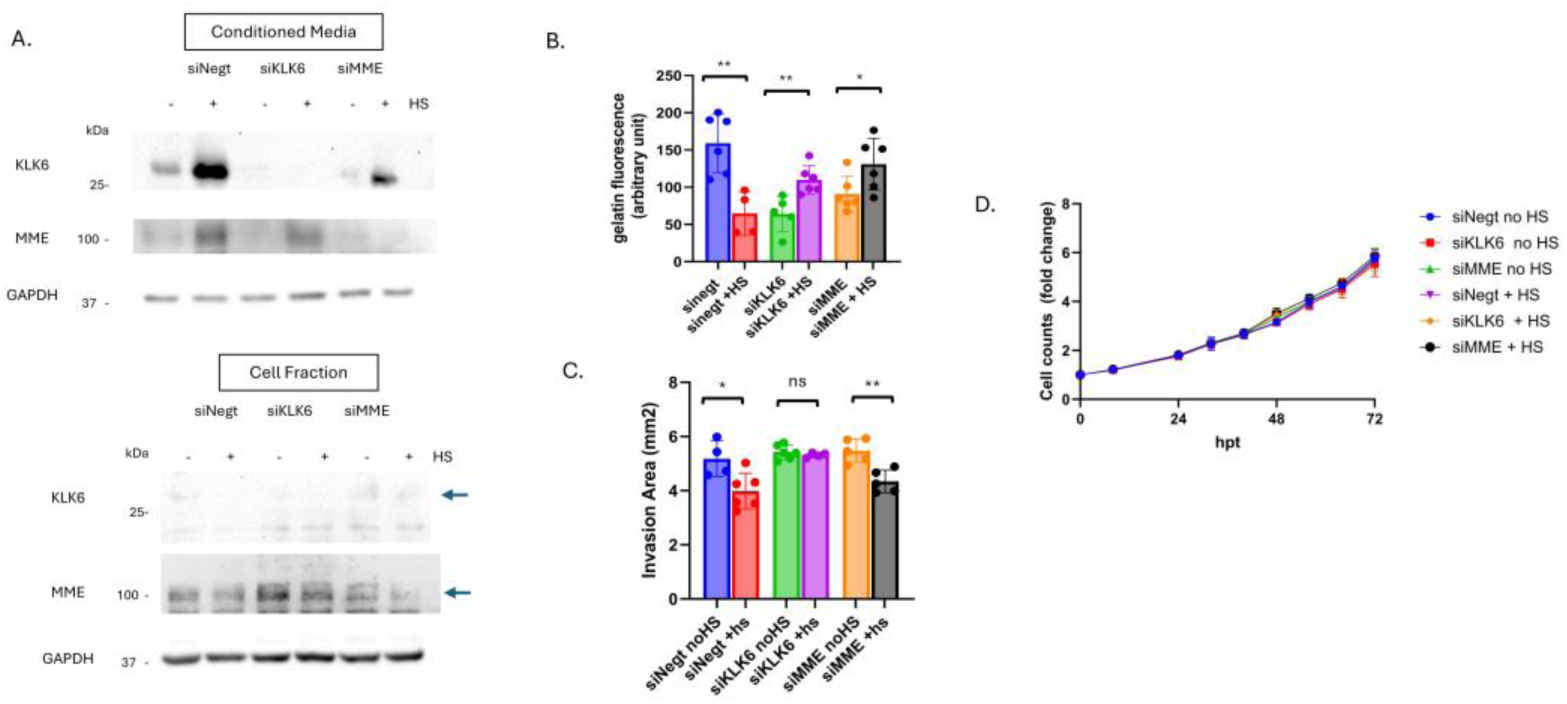
Stress-induced KLK6 impacts MMP-2 activity and cellular invasion. (**A**) Immunoblot showing siRNA knockdown of KLK6 and MME in PC3 cell fraction and conditioned media 72hr post transfection and 24hr post stress n=3 (**B**) Gelatin DQ assay measuring MMP-2 activity in CM from siRNA samples -/+ stress. (**C**) Matrigel drop invasion assays with untreated PC3 cells incubated with CM from siRNA KD plates -/+ stress. Invasion was measured over 4 days. (**D**) Proliferation assay with untreated PC3 cells incubated with CM from untreated and stressed siRNA samples. A minimum of 2 individual biological replicates were performed for the above experiments, and the results were combined. Statistical significance was determined by Welch’s student t-test. * p<0.05, **<0.01.

We next examined effects of MME or KLK6 knockdown on MMP-2 activity. Gelatin DQ assays showed that knockdown of either MME or KLK6 results in lower MMP-2 activity in non-stressed cells. When subjected to stress, knockdown of either MME or KLK6 rescued MMP-2 activity (Figure 8B). To examine invasion, CM from untreated and stressed knockdown samples were used in Matrigel drop invasion assays (Figure 8C). siNegt CM mimicked what we previously observed: CM from stressed cells significantly decreased cellular invasion rates. However, CM lacking KLK6 rescued stress-induced decreases in cell invasion. In contrast, CM from stressed cells lacking MME did not rescue decreased invasion, likely because KLK6 was still present in CM from these samples and expression was increased upon stress. Therefore, only when KLK6 was present in CM (in siNegt and siMME transfection samples) did stress lead to decreased invasion. This indicates the importance of KLK6 in this process; a role that is reliant on stress-induced expression as well as its subsequent inhibition of MMP-2. Once KLK6 was removed or inhibited, stress did not result in decreased MMP activity nor cell invasion. To ensure invasion rates were not affected by changes in proliferation we conducted growth assays with CM from non-stressed and stressed knockdown cells applied to untreated PC3 cells (Figure 8D). Cell counts were monitored over 72 hours. Proliferation was not altered by CM from untreated or stressed knockdown samples. Together, these findings identify KLK6 as a protease that decreases MMP-2 activity and cell invasion in response to stress.

## 4. Discussion

This study reveals a novel regulatory network governing cell migration and invasion in PCa. While eHSP90α has previously been associated with increased MMP-2 activity, we expanded on the unique finding that in the context of stress-induced eHSP90α, MMP-2 activity was decreased. This was discovered to be due to stress-induced expression of the serine protease KLK6, when at higher extracellular levels resulted in decreased MMP-2 activity and decreased cell invasion. Inhibition or knockdown of KLK6 resulted in rescued activity and invasion. This rescue was dependent on HSP90α, as it could not be seen in HSP90α KO cells (Figure 9).

**Figure 9.**
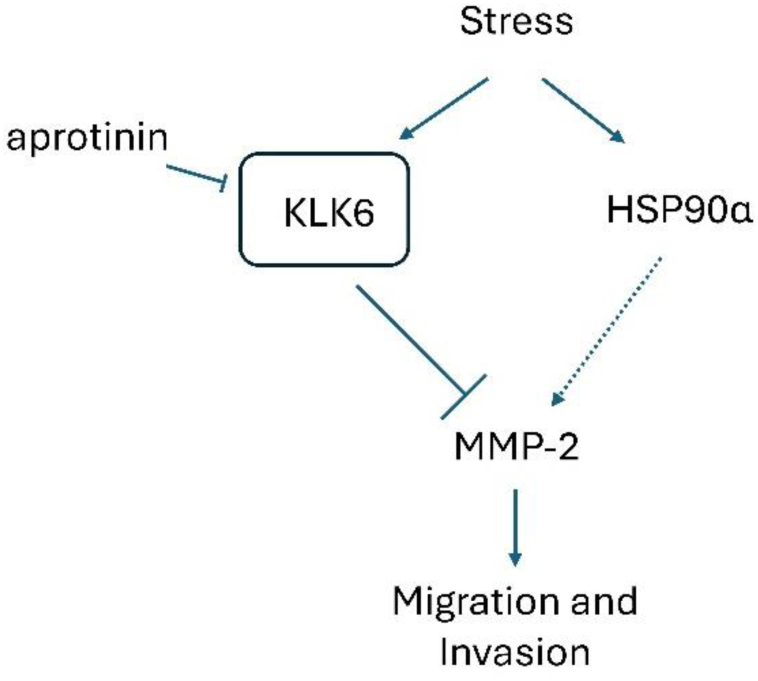
Model of regulatory stress-response proteins driving MMP-2-dependent PCa metastatic behavior. Stress induces increased levels of extracellular regulatory proteins: chaperone HSP90α and serine protease KLK6. This also leads to a decrease in MMP-2 activity and cell invasion. Addition of aprotinin, a serine protease inhibitor, or knockdown of KLK6 is enough to rescue this effect and increase MMP activity and cell invasion in PCa cells. This rescue is dependent on HSP90α only once KLK-6 is inhibited.

Our study supports findings that eHSP90α is linked to MMP-2 activity. Stress-induced increased MMP-2 activity was only observed once KLK-6 was inhibited in the presence of HSP90α. It is likely that eHSP90α is maintaining MMP-2 activity through its chaperone function. Baker-Williams et al. show an important mechanism involving TIMP2 and AHA1 as co-chaperones of eHSP90α where the two act as a molecular switch to inhibit or induce eHSP90α MMP-2 activation [75]. Another study demonstrated an ATP-independent function of eHSP90α binding to and stabilizing MMP-2 or protecting it from degradation [76]. If these mechanisms are at play in our system, they may explain the role of eHSP90α in rescued MMP-2 activity once KLK-6 is inhibited or removed. Co-chaperone proteins of eHSP90α will be important to consider in future mechanistic studies with PCa, especially as the above studies do not examine MMP activity in the context of cell stress.

The role of KLK6 in cancer is context dependent. Increased expression is associated with poor prognosis in several cancer types including ovarian, colon, gastric, and pancreatic [77–85]. In breast cancer cells, KLK6 has been shown to act as a tumor suppressor [86]. A separate report found that increased levels of KLK6 in serum from patients with invasive breast cancer, compared to healthy controls, suggesting a role in disease progression and poor prognosis [87]. However, in head and neck squamous cell carcinoma, higher levels of KLK6 were associated with favorable patient survival. In head and neck squamous cell cancer (HNSCC) cancer, a tumor suppressor function is supported by showing that knockdown of KLK6 led to increased proliferation, migration, invasion, epithelial-to-mesenchymal transition markers and survival post radiation [88]. Our findings align with those in found in HNSCC and breast cancer as it demonstrates that higher KLK-6 expression decreases MMP-2 activity and invasion potential in PCa.

As for linking KLK6 to MMP activity, this has mainly been looked at in neuronal cells. In vitro studies show KLK6 can cleave pro-MMP-2 and pro-MMP-14. However, it does not cleave MMPs at the same canonical activating site as Plasmin [89, 90]. It is suggested that removing part of MMP-2 pro-domain may prime or partially activate it. In our case, it may render it inactive. If MMP-2 is further processed and degraded when KLK6 is upregulated upon stress this could also lead to decreased activity.

While we do not show a strong effect on cell invasion with MME knockdown, it may still contribute to a smaller degree, or its role could be masked by other stress response proteins such as KLK6 or eHSP90α. MME influences multiple cellular pathways through the cleavage of extracellular activating peptides. It indirectly affects various transduction pathways in tumor development [91, 92]. It’s role in cancer biology is also context dependent. Decreased MME levels are associated with disease progression in many solid tumors such as breast, cervix, ovary, and lung [93–98]. Alternatively, increased levels are associated with advanced melanoma, leukemia, and colon cancers [99–104]. In PCa, MME is associated with decreased cell migration and disease progression [105–107]. We demonstrate that knockdown of MME in PC3 cells, however, resulted in lower invasion when cells were treated with HS CM. This could be due to KLK6 increase. While the CM of MME knockdown cells that were non-stressed had little impact on invasion, it is likely because there was very little MME expression to start with. MME is usually a membrane bound peptidase, yet it can be secreted in extracellular vesicles or as a soluble protein [108–112]. It is not clear in our study which extracellular form MME is secreted by PCa cells.

HSP90’s role in cancer biology has been extensively studied and historically therapeutics have been toxic as it is vital in healthy cells as well. However, eHSP90α shines a new light on therapeutic targeting strategies [17, 18, 113]. Utilizing novel HSP90 interacting therapeutics pioneered by our group [114], as well as cell impermeable therapeutics such as antibodies, beads, or non-permeable drug analogs could greatly improve outcomes while avoiding toxicity. Our data indicates KLK6 expression is favorable in PCa, but this is contingent upon eHSP90α status. eHSP90α may be an activator, protector and/or chaperone of MMP-2 [19–22, 24, 75]. Therefore, targeting eHSP90α would be ideal while KLK6 status would be informative. A notable challenge in targeting eHSP90α is isoform specificity. HSP90α and HSP90β share 86% homology at the amino acid level. If a potential therapeutic could be maintained outside the cell this would help immensely in reducing toxicity and targeting only eHSP90α, as eHSP90β does not seem to play a role in ECM invasion [10, 115, 116]. In addition to expression, localization, isoform specificity, markers of cellular stress would also be important to consider.

This study reports the first established PCa HSP90α KO cells. Previous reports of HSP90α knockout in human cell lines include HEK-293T, A549 (lung cancer), MDA-MB-231 (breast cancer) and HT1080 (fibrosarcoma) [116–120]. Knockout mice and MEFs have also been reported [121–123]. While it may seem counterintuitive that PCa KO cells have increased extracellular MMP-2 activity and the cells themselves have decreased cell motility, studies in BrCa cells also report decreased cell motility in HSP90α KO cells [116]. These studies do not specifically examine MMP-2 activity.

Extracellular proteins are ideal biomarkers, as well as important therapeutic targets. Clinical diagnostics utilizing liquid biopsy have powerful implications for eHSP90α and KLK6. Serum levels of KLK6 in ovarian and invasive BrCa patients were significantly higher than non-cancer controls [78, 87]. This may vary in PCa depending on stress response proteins or advancement of disease, and/or treatment regimes. HSP90α has been looked at in serum from many different cancer types over the past two decades [17]. It is significantly higher in liver, lung, melanoma, hepatocellular carcinoma, cervical, prostate, gastric and acute myeloid leukemia cancer patients compared to healthy controls [124–134]. While not currently accepted as a standard clinical diagnostic it holds high potential to add to current methods used for risk stratification and tailored care for patients.

## Author Contributions

Conceptualization, K.L.O., J.Z., R.B.; methodology, K.L.O., J.Z.; validation, K.L.O., J.Z.; formal analysis, K.L.O.; investigation, K.L.O., J.Z.; resources, R.B.; data curation, K.L.O., J.Z.; writing—original draft preparation, K.L.O., R.B.; writing—review and editing, K.L.O., J.Z., R.B..; visualization, K.L.O., J.Z.; supervision, K.L.O., R.B.; funding acquisition, K.L.O., R.B.; All authors have read and agreed to the published version of the manuscript.

## Funding

This research was funded by the National Institutes of Health to K.L.O. [R01CA276846-S1] and to R.B. [R01CA276846]

## Data Availability Statement

All data are contained within the article or Supplementary Materials.

## Acknowledgments

We would like to thank Dr. Grinu Mathew, Dr. Abdalla Maher, Dr. Fangfang Qiao, Dr. Lianmei Zhao, Weining Chen, Jaclyn Knibbe-Hollinger, and Rachel Rhatigan for helpful discussions and advice on experimental protocols and approaches. We thank Dr. David Kelly and Pei Xian Chen for help with use of the Incucyte S3 as well as data analysis protocols. We are grateful to Dr. Suzanne L. Topalian for the generous gift of HPV transformed primary cell lines: 1532NPTX and 1532CPTX. We also thank Dr. Ken-Ichi Takemaru for the gift of the CRISPR plasmid: pSpCas9(BB)-2A-Blast.

## Conflicts of Interest

The authors declare no conflicts of interest in this study.

The following abbreviations are used in this manuscript:

## Abbreviations

PCa: Prostate cancer
MMP-2: matrix metalloproteinase-2
eHSP90α: Extracellular heat shock protein 90α
KO: knockout
KD: knockdown
KLK6: kallikrein-related peptidase 6
HSP90: heat shock protein 90
MMP: matrix metalloproteases
ECM: Extracellular matrix
BrCa: Breast cancer
ATCC: American Type Culture Collection
rEGF: recombinant epidermal growth factor
BPE: bovine pituitary extract
HS: heat shock
SFM: serum free media
CF: cell fraction
CM: conditioned media
DQ: dequenching
Hr: hour
PCR: Polymerase chain reaction
DAPI: 4′,6-diamidino-2-phenylindole
PCR: Polymerase chain reaction
AR: Androgen receptor
iHSP90α: Intracellular heat shock protein 90α
PPI: protease/phosphatase inhibitors
EV: Empty vector control
PI: Protease inhibitors
PhI: Phosphatase inhibitors
MME: membrane metallo-endopepitdase
HNSCC: head and neck squamous cell carcinoma

## Notes

### Competing Interest Statement

The authors have declared no competing interest.

### Summary of Updates

No major differences. Supplemental files updated. File type revised.

