## Supplemental Figures 1-8 for "HSP90α and KLK6 Co-Regulate Stress-Induced Prostate Cancer Cell Motility"

### Slide 1
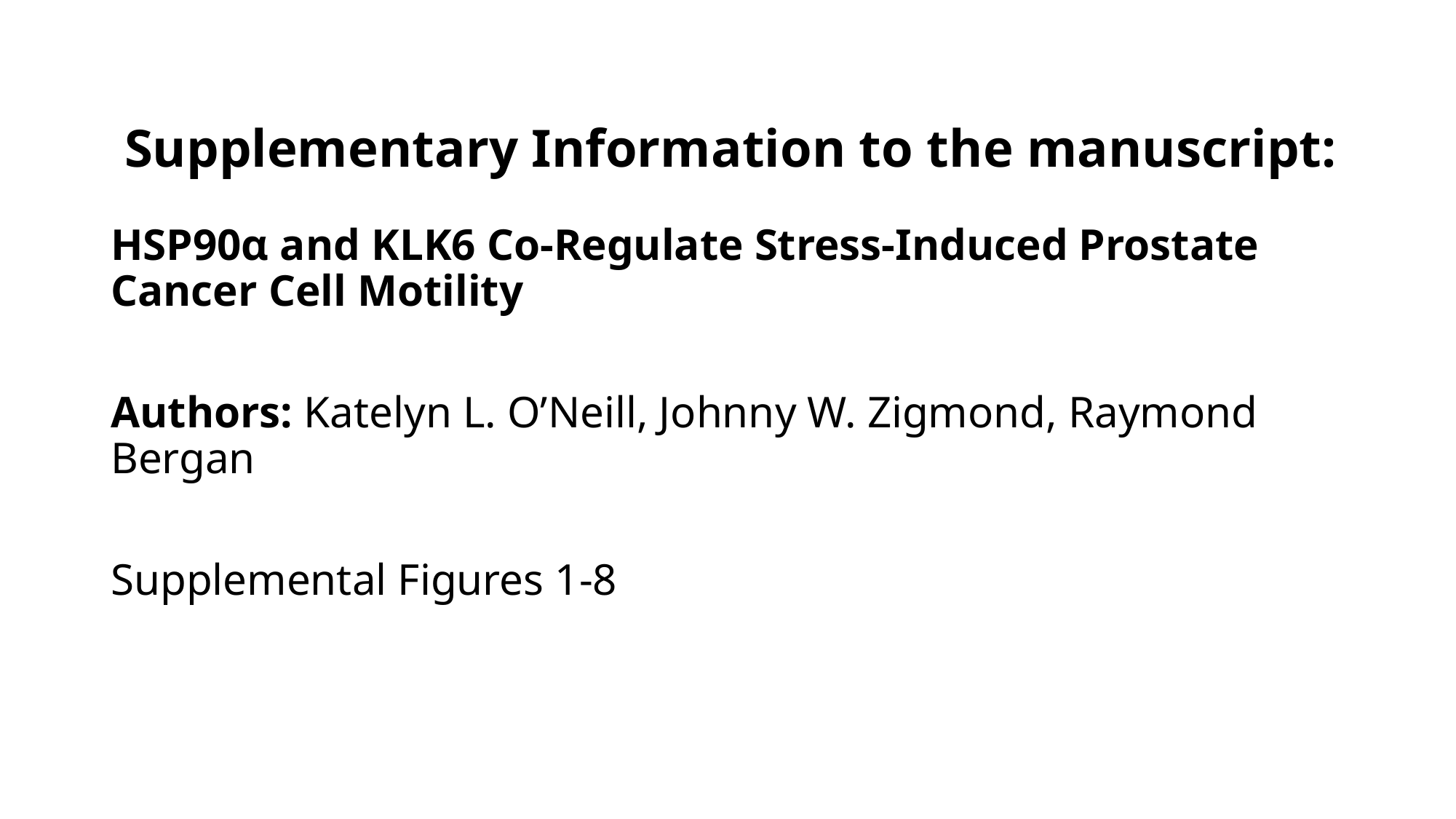

#### Supplementary Information to the manuscript:
HSP90α and KLK6 Co-Regulate Stress-Induced Prostate Cancer Cell Motility
Authors: Katelyn L. O’Neill, Johnny W. Zigmond, Raymond Bergan
Supplemental Figures 1-8

### Slide 2
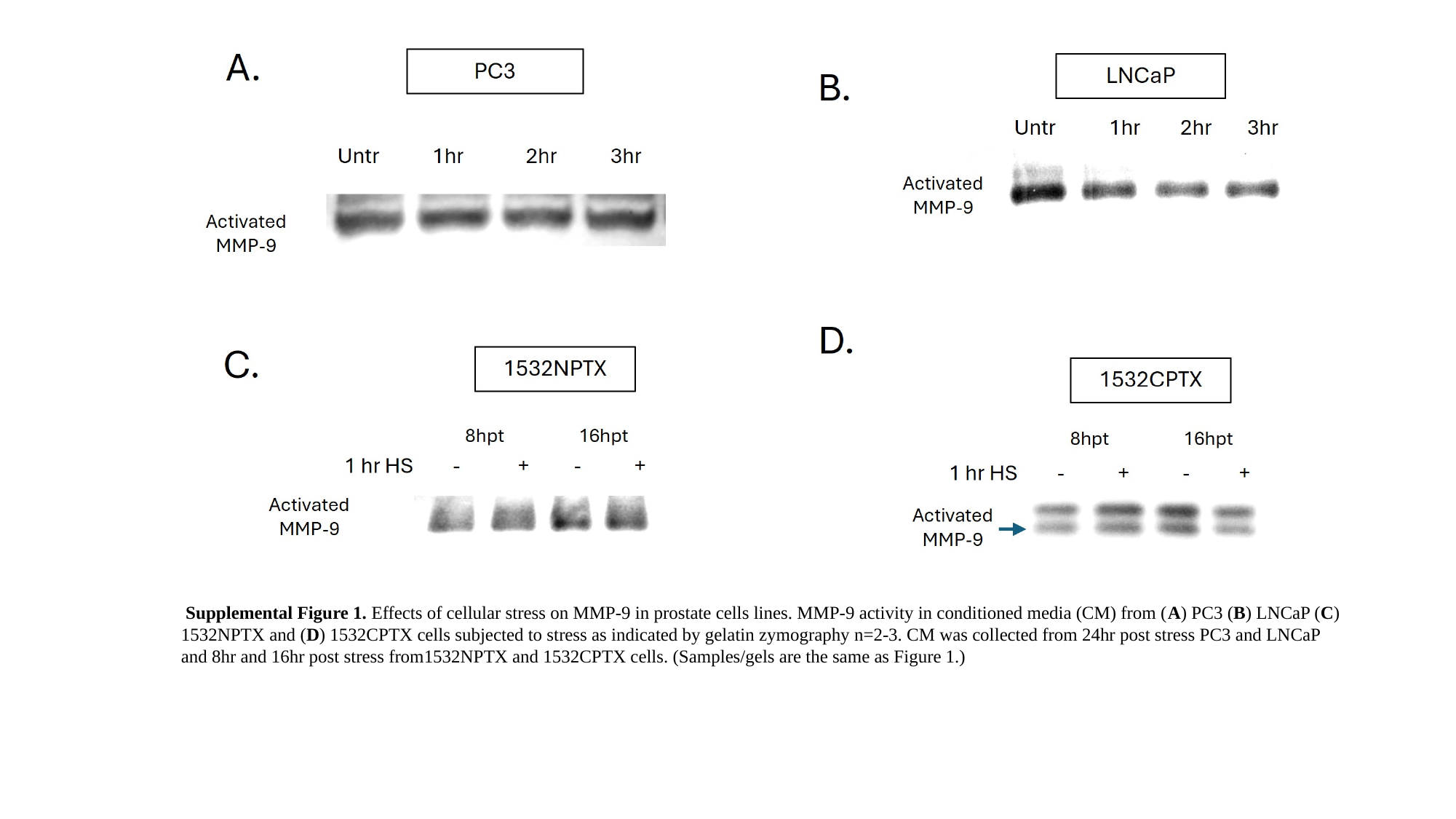

Supplemental Figure 1. Effects of cellular stress on MMP-9 in prostate cells lines. MMP-9 activity in conditioned media (CM) from (A) PC3 (B) LNCaP (C) 1532NPTX and (D) 1532CPTX cells subjected to stress as indicated by gelatin zymography n=2-3. CM was collected from 24hr post stress PC3 and LNCaP and 8hr and 16hr post stress from1532NPTX and 1532CPTX cells. (Samples/gels are the same as Figure 1.)​
​

### Slide 3
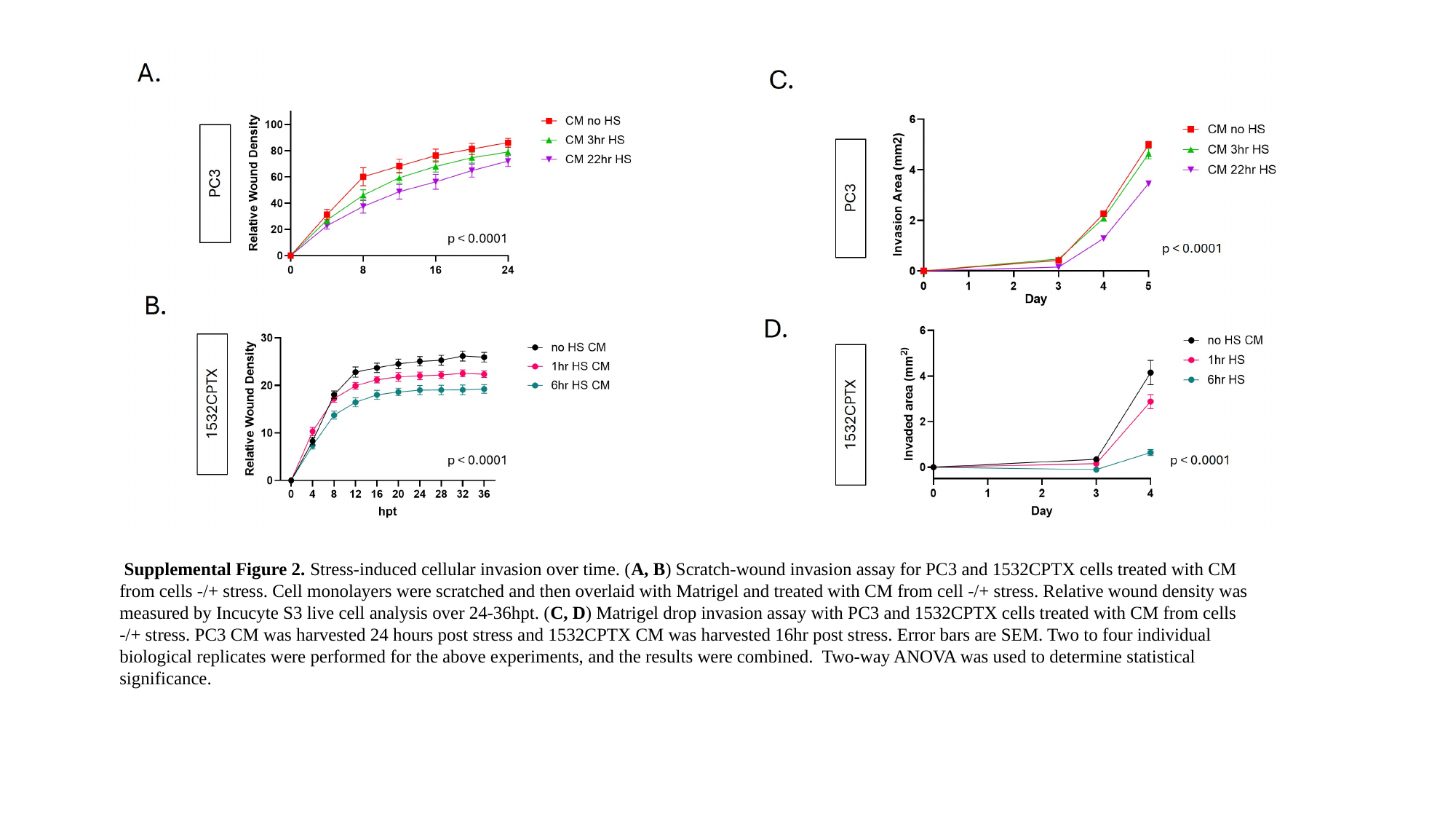

Supplemental Figure 2. Stress-induced cellular invasion over time. (A, B) Scratch-wound invasion assay for PC3 and 1532CPTX cells treated with CM from cells -/+ stress. Cell monolayers were scratched and then overlaid with Matrigel and treated with CM from cell -/+ stress. Relative wound density was measured by Incucyte S3 live cell analysis over 24-36hpt. (C, D) Matrigel drop invasion assay with PC3 and 1532CPTX cells treated with CM from cells -/+ stress. PC3 CM was harvested 24 hours post stress and 1532CPTX CM was harvested 16hr post stress. Error bars are SEM. Two to four individual biological replicates were performed for the above experiments, and the results were combined. Two-way ANOVA was used to determine statistical significance.

### Slide 4
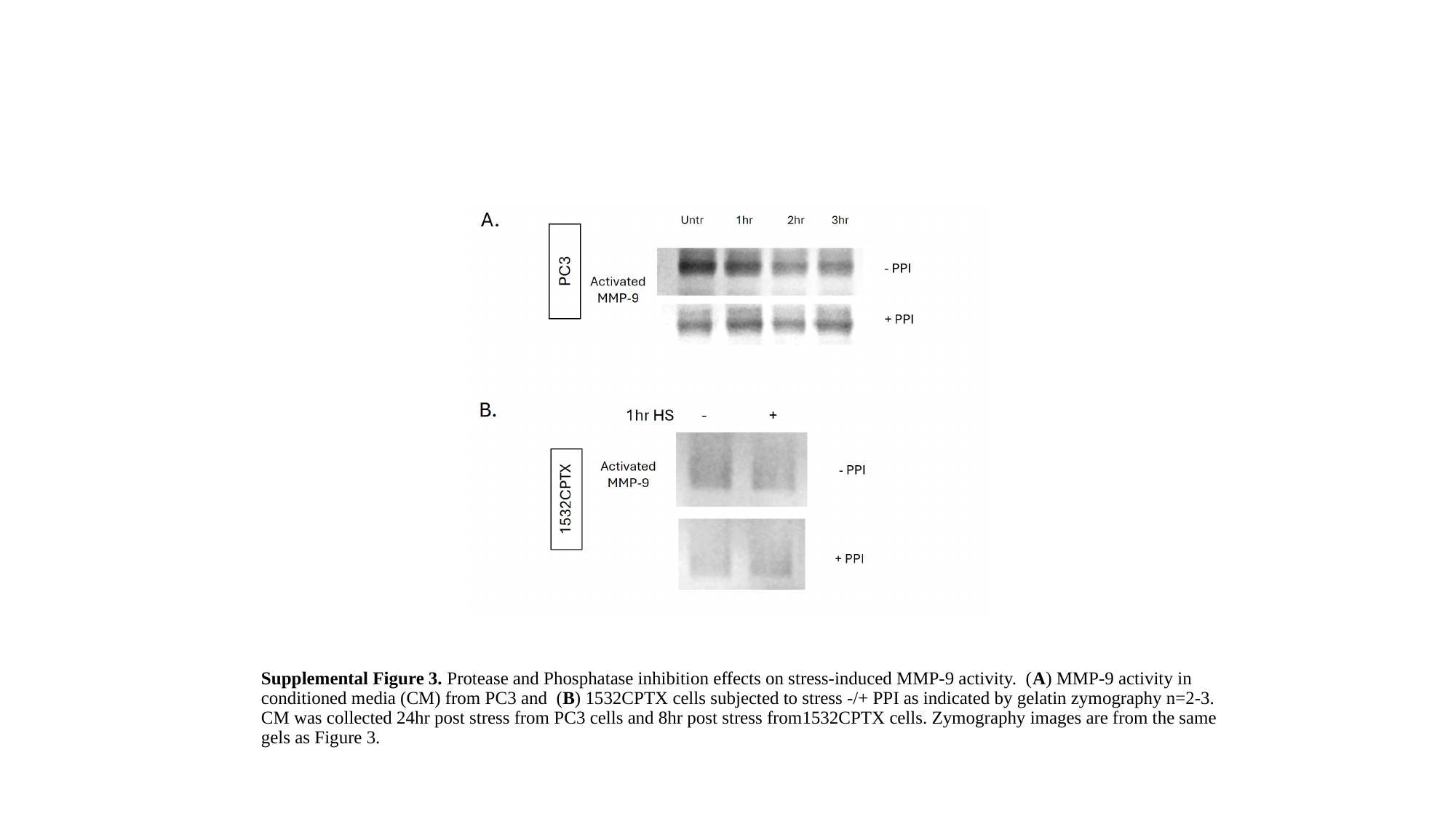

Supplemental Figure 3. Protease and Phosphatase inhibition effects on stress-induced MMP-9 activity. (A) MMP-9 activity in conditioned media (CM) from PC3 and (B) 1532CPTX cells subjected to stress -/+ PPI as indicated by gelatin zymography n=2-3. CM was collected 24hr post stress from PC3 cells and 8hr post stress from1532CPTX cells. Zymography images are from the same gels as Figure 3.

### Slide 5
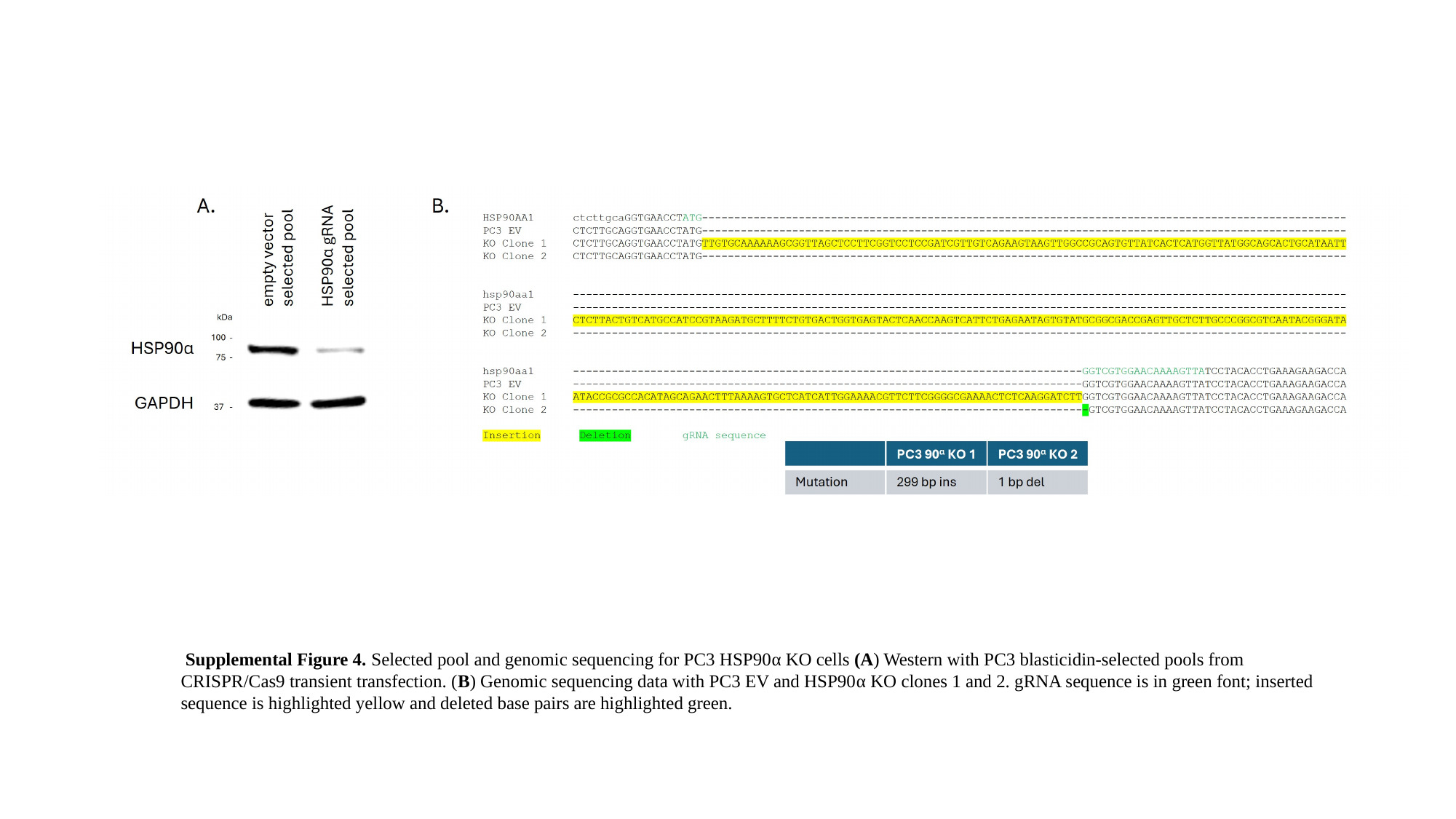

Supplemental Figure 4. Selected pool and genomic sequencing for PC3 HSP90α KO cells (A) Western with PC3 blasticidin-selected pools from CRISPR/Cas9 transient transfection. (B) Genomic sequencing data with PC3 EV and HSP90α KO clones 1 and 2. gRNA sequence is in green font; inserted sequence is highlighted yellow and deleted base pairs are highlighted green.

### Slide 6
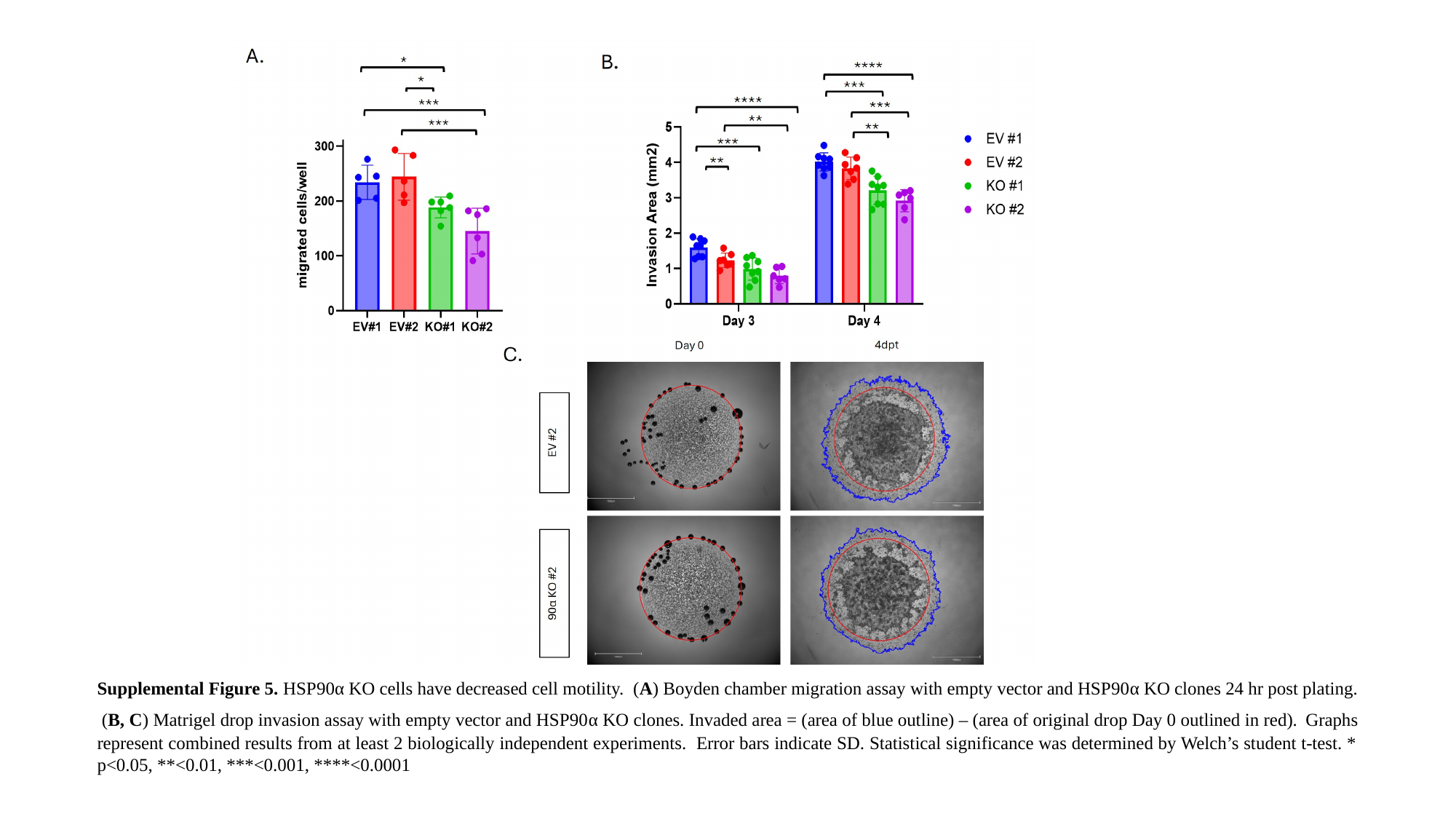

Supplemental Figure 5. HSP90α KO cells have decreased cell motility. (A) Boyden chamber migration assay with empty vector and HSP90α KO clones 24 hr post plating. (B, C) Matrigel drop invasion assay with empty vector and HSP90α KO clones. Invaded area = (area of blue outline) – (area of original drop Day 0 outlined in red). Graphs represent combined results from at least 2 biologically independent experiments. Error bars indicate SD. Statistical significance was determined by Welch’s student t-test. * p<0.05, **<0.01, ***<0.001, ****<0.0001

### Slide 7
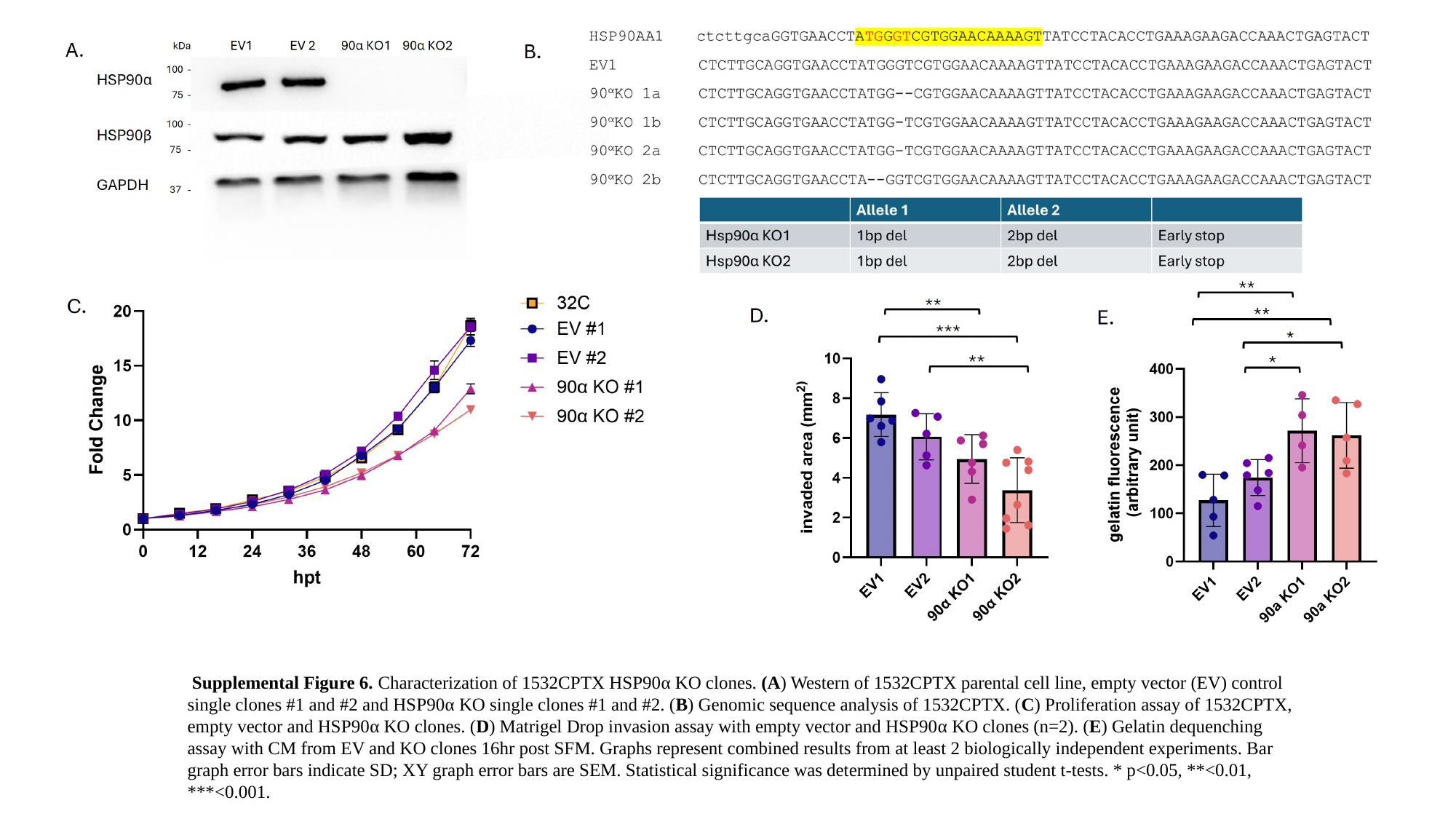

Supplemental Figure 6. Characterization of 1532CPTX HSP90α KO clones. (A) Western of 1532CPTX parental cell line, empty vector (EV) control single clones #1 and #2 and HSP90α KO single clones #1 and #2. (B) Genomic sequence analysis of 1532CPTX. (C) Proliferation assay of 1532CPTX, empty vector and HSP90α KO clones. (D) Matrigel Drop invasion assay with empty vector and HSP90α KO clones (n=2). (E) Gelatin dequenching assay with CM from EV and KO clones 16hr post SFM. Graphs represent combined results from at least 2 biologically independent experiments. Bar graph error bars indicate SD; XY graph error bars are SEM. Statistical significance was determined by unpaired student t-tests. * p<0.05, **<0.01, ***<0.001.

### Slide 8
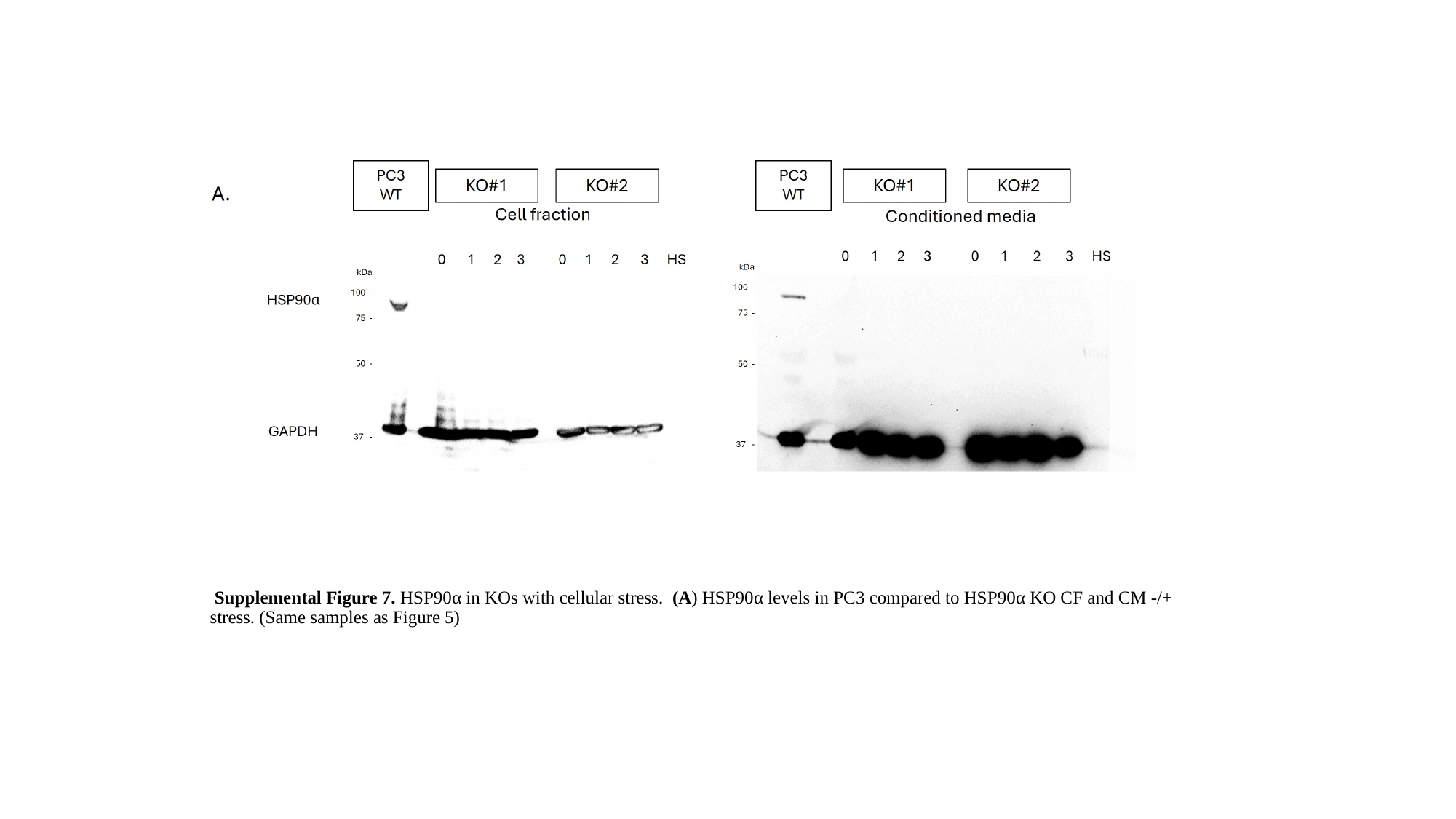

Supplemental Figure 7. HSP90α in KOs with cellular stress. (A) HSP90α levels in PC3 compared to HSP90α KO CF and CM -/+ stress. (Same samples as Figure 5)

### Slide 9
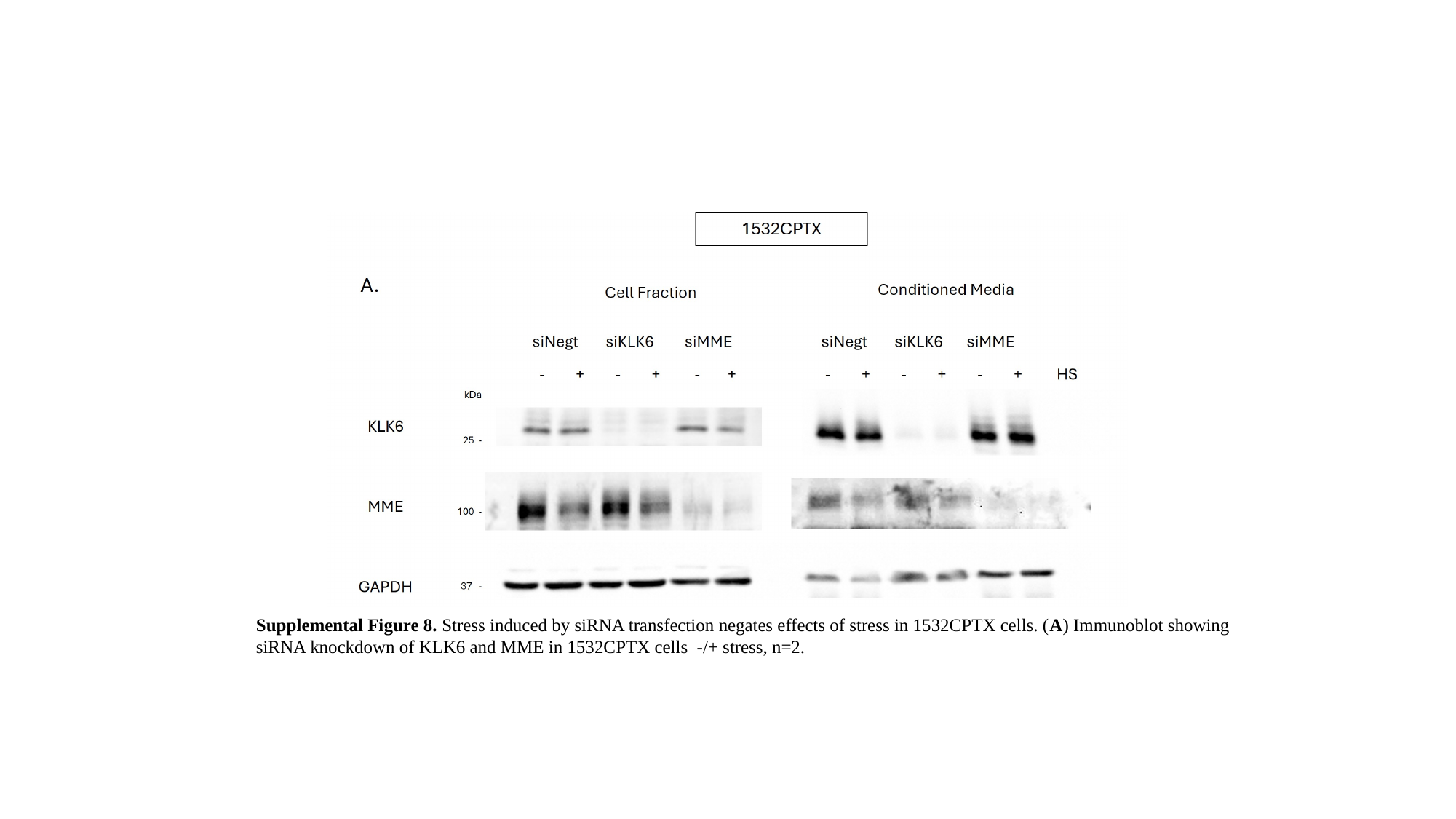

Supplemental Figure 8. Stress induced by siRNA transfection negates effects of stress in 1532CPTX cells. (A) Immunoblot showing siRNA knockdown of KLK6 and MME in 1532CPTX cells -/+ stress, n=2.
